# Excitation/Inhibition Balance in People at Clinical High Risk for Psychosis: A Functional MRI Study

**DOI:** 10.64898/2026.11.24.768831

**Authors:** Zilin Li, Ioana Varvari, Alexis E. Cullen, George Gifford, Paola Dazzan, Stefania Tognin, Paolo Fusar-Poli, Gemma Modinos, Rene Kahn, Erika van Hell, Arija Maat, Lieuwe de Haan, Benedicto Crespo-Facorro, Birte Glenthøj, Stephen Lawrie, Oliver Gruber, Thérèse van Amelsvoort, Celso Arango, Tilo Kircher, Barnaby Nelson, Silvana Galderisi, Rodrigo A. Bressan, Jun Soo Kwon, Mark Weiser, Romina Mizrahi, Gabriele Sachs, Matthias Kirschner, Anke Maatz, Samuel R. Knight, Timothy Lawn, Ofer Pasternak, Dominic Oliver, Robert McCutcheon, Philip McGuire, the PSYSCAN Consortium

## Abstract

Excitation/inhibition (E/I) imbalance is implicated in the pathophysiology of psychosis, but the extent to which this is evident before the onset of illness is unclear. Here, we investigated E/I-related brain dynamics using the Hurst exponent (HE) derived from resting-state MRI in two longitudinal cohorts of individuals at clinical high risk for psychosis (CHR-P): PSYSCAN (190 CHR-P individuals and 115 controls) and Accelerating Medicines Partnership® Schizophrenia (826 CHR-P individuals and 241 controls). Both cohorts showed lower whole-brain HE in CHR-P individuals than in controls at baseline, with the most prominent differences observed in the somatomotor network and thalamus. In both cohorts, normative receptor distributions significantly predicted the spatial pattern of HE differences, with consistent contributions from norepinephrine, serotonin, and opioid receptors. In PSYSCAN, 21 CHR-P individuals transitioned to psychosis during follow-up. There was preliminary evidence that subsequent transition to psychosis was associated with a longitudinal reduction in HE in the right inferior parietal cortex and that baseline regional HE showed above-chance prediction of subsequent transition at the individual level. Together, these findings suggest that vulnerability to psychosis is associated with reduced HE, consistent with less persistent brain dynamics and increased excitability.

## 1. Introduction

Genetic, pharmacological, and post-mortem evidence suggests that synaptic dysfunction plays a critical role in the pathophysiology of psychosis.^1–5^ This appears to involve the interactions between excitatory glutamatergic and inhibitory GABAergic neurons. The excitation/inhibition (E/I) balance between these groups of neurons is essential for efficient information processing within and between neural circuits.^6,7^ One hypothesis is that psychosis is associated with N-methyl-D-aspartate receptor (NMDAR) hypofunction on GABAergic interneurons, which would reduce inhibitory control of pyramidal neurons, resulting in disinhibition and cortical hyperactivity.^8,9^ These changes may ultimately lead to large-scale neural network dysfunction and the emergence of psychotic symptoms.^8–10^

E/I balance cannot be measured at the neuronal level in living humans. Recent work, however, has suggested that functional magnetic resonance imaging (fMRI), electroencephalography (EEG) and magnetic resonance spectroscopy (MRS) may be able to quantify proxy measures of underlying E/I balance. The Hurst exponent (HE) measures long-range temporal autocorrelations and, when applied to fMRI time-series, has been proposed as a reliable index for E/I balance based on converging in silico, in vivo, and human evidence.^11–13^ A shift towards excitation is thought to reduce the temporal persistence of neural activity, leading to weaker long-range autocorrelations and therefore a lower HE.^12^ HE has also been associated with parvalbumin (PV) mRNA expression in humans and PV-positive cell density in mice,^14^ suggesting a potential link to PV interneurons, whose dysfunction has been implicated in E/I imbalance in schizophrenia.^15,16^

fMRI and EEG studies suggest that proxy measures of E/I balance are significantly altered in patients with psychosis.^17–19^ However, because these data were collected from participants in whom psychosis was well established, it is unclear whether these changes are related to the disorder or reflect risk factors for its development. This issue can be addressed by assessing E/I balance in people at clinical high risk for psychosis (CHR-P), who have an increased vulnerability to psychosis, but have not yet developed the disorder.^20^

The present study aimed to investigate E/I-related brain dynamics in people at CHR-P using resting-state fMRI (rsfMRI). If E/I-related alterations are associated with an increased vulnerability to psychosis, then these would be expected to be evident in this group. If these alterations are associated with subsequent transition to psychosis, they may be more pronounced or show different trajectories in the minority of CHR-P individuals who later developed psychosis. We compared HE between CHR-P individuals and healthy controls (HCs) and examined its longitudinal trajectories in two independent cohorts, PSYSCAN and the Accelerating Medicines Partnership® Schizophrenia (AMP® SCZ). In PSYSCAN, we additionally compared CHR-P individuals who transitioned to psychosis (CHR-T) with those who did not (CHR-NT). We characterized the spatial patterns of HE alterations at the whole-brain, network, and regional levels and further examined their associations with potential neurobiological substrates. We hypothesised that HE would be lower in CHR-P individuals than in HCs and that HE alterations would be associated with subsequent transition to psychosis.

## 2. Methods

### Study design

#### Cohort 1

CHR-P individuals and HCs were recruited between July 2016 and December 2019 from PSYSCAN,^21,22^ a multi-centre prospective study. Participants with imaging data were enrolled from 12 sites: London, Amsterdam, Maastricht, Madrid, Naples, Melbourne, Seoul, Hong Kong, Toronto, São Paulo, Edinburgh, Utrecht.

#### Cohort 2

For independent validation, CHR-P individuals and HCs were recruited between June 2022 and January 2025 from another multi-centre prospective study, Accelerating Medicines Partnership® Schizophrenia (AMP® SCZ).^23^ The dataset used in this study was downloaded from the NIMH Data Archive (data release 3.0 doi: 10.15154/81xe-k706). Participants with imaging data were enrolled from 38 sites in the United States, Canada, the United Kingdom, Spain, Italy, Switzerland, Netherlands, Germany, Denmark, Australia, Singapore, South Korea, Chile.

All participants provided written informed consent under institutional review boards approved procedures. Details of both cohorts have been published elsewhere.^21–23^

#### Participants

In PSYSCAN, CHR-P status was defined using either the Comprehensive Assessment of At-Risk Mental States (CAARMS)^24^ or the Schizophrenia Proneness Instrument (SPI-A).^25^ Transition to psychosis in CHR-P individuals was assessed at 3, 6, 12, 18, and 24 months. Follow-up assessments were extended beyond 24 months when feasible or necessary to maximize data completeness, with additional data obtained through face-to-face assessments or electronic health records (EHRs). Transition to psychosis was defined either psychometrically using CAARMS criteria or clinically using Diagnostic and Statistical Manual of Mental Disorders (DSM)- or the International Classification of Diseases (ICD)-defined diagnoses or accepted referrals to early intervention services recorded in EHRs.^22^

In AMP SCZ, CHR-P status was defined using the Positive SYmptoms and Diagnostic Criteria for the CAARMS Harmonized with the SIPS (PSYCHS).^26^ Transition outcome data in the AMP SCZ were not available as clinical follow-up was still ongoing. Details on participant eligibility are provided in **eMethods 1**.

#### Clinical measures

In PSYSCAN, positive/negative symptoms were assessed using the CAARMS.^24^ Global functioning was measured using the Social and Occupational Functioning Assessment Scale (SOFAS).^27^

In AMP SCZ, positive symptoms were assessed using the PSYCHS.^26^ Negative symptoms were assessed using the Negative Symptom Inventory–Psychosis Risk (NSI-PR).^28^ Functioning was measured with SOFAS.^27^

#### rsfMRI data acquisition and preprocessing

rsfMRI data were acquired at baseline, 6 months, and 12 months in PSYSCAN, and at baseline and 2 months in AMP SCZ using 3T scanners with gradient-echo echo-planar imaging. Preprocessing followed prior work validating HE estimation^12^: core preprocessing with fMRIPrep v23.2.0,^29^ followed by denoising using the wavelet-based time-series despiking^30^ and 13 parameter nuisance regression (6 motion parameters, their first derivatives, and cerebrospinal fluid signal). Mean time series were extracted from 426 parcels (66 subcortical regions) using the Human Connectome Project Extended (HCPex) atlas.^31^ Imaging parameters, preprocessing and quality-check details are in **eMethods 2**.

#### Hurst exponent (HE) calculation

Consistent with prior HE work,^12^ participants with a mean framewise displacement (meanFD) > 0.4 mm were excluded. Sites with < 10 participants were also excluded. Regional HE was estimated using a discrete wavelet transform and a fractionally integrated process model with maximum likelihood estimation.^12^

To address scanner-related variability, PSYSCAN sites with travelling-head data were corrected using site-specific offsets and then treated as a single mega-site. The remaining sites were harmonized with this mega-site using neuroComBat.^32^ In AMP SCZ, where travelling-head data were unavailable, all sites were harmonized with neuroComBat. In both cohorts, neuroComBat included site as batch variable and age, sex, phenotype/transition status, MeanFD as covariates.

Whole-brain HE was calculated as the mean across all parcels. Network-level analyses were performed using the Yeo seven-network parcellation,^33^ including the Visual (VN), Somatomotor (SMN), Dorsal Attention (DAN), Salience (SN), Limbic (LN), Frontoparietal (FPN), and Default Mode (DMN) networks (**eFigure 2**). Details of HE estimation, site harmonization, and network mapping are in **eMethods 3**.

### Statistical analysis

#### Group and Transition-Related Differences in HE

Group differences in age, symptoms, and functioning were assessed using Welch’s t-tests, and sex, antipsychotic use using chi-square tests.

HE differences were examined at the global, network-wise and parcel-wise levels. Ordinary least squares (OLS) regression models were used to examine baseline HE differences between CHR-P individuals and HCs in both datasets, and between CHR-T and CHR-NT individuals in PSYSCAN only. The model was: HE = β₀ + β₁Group + β₂Age + β₃Sex + β₄MeanFD + ε.

To quantify the cross-cohort spatial correspondence of regional case-control effects, Pearson correlations were calculated between parcel-wise phenotype β₁ maps in PSYSCAN and AMP SCZ, separately for cortical and subcortical regions. Statistical significance was assessed using 10,000 subject-level phenotype-label permutations.

Linear mixed-effects models were used to investigate longitudinal alterations in HE. Group, time, and their interaction were included as fixed effects, with time treated as a categorical variable, and subject ID was included as a random intercept. The model was: HE = β_0 +_ β_1G_roup + β_2T_ime + β_3(_Group×Time) + β_4A_ge+β_5S_ex + β_6M_eanFD + b_ID +_ ε.

Multiple comparisons across parcels or networks were false discovery rate (FDR) corrected (q < 0.05). The effect of antipsychotic exposure was assessed only in PSYSCAN by comparing whole-brain HE between participants with and without antipsychotic treatment using Welch’s t-test, as no CHR-P subjects in AMP SCZ were receiving antipsychotic medication.

#### Individual-level prediction of psychosis transition

To complement the group-level analyses, we examined whether baseline parcel-wise HE could predict subsequent transition to psychosis at the individual level in the PSYSCAN CHR-P sample. A prediction model was developed using elastic net-regularized Cox proportional hazards regression within a repeated nested cross-validation framework. Performance was evaluated using Harrell’s C-index, balanced accuracy (BAC), sensitivity, specificity and Brier score. Details of model development are provided in **eMethods 4**.

#### Biological correlates of HE phenotype effect

To explore potential biological correlates of regional HE alterations, we examined their spatial correspondence with cortical structural differences and normative neurotransmitter receptor/transporter distributions. Cortical thickness (CT) was calculated using FreeSurfer^34^ and we assessed the spatial correspondence between phenotype-associated effects in HE and CT using Pearson correlation. Nineteen receptor/transporter density maps were obtained from the Hansen atlas^35^ and used to predict case-control HE differences using partial least-squares regression (PLSR) with nested cross-validation. Model performance was evaluated using cross-validated R² and Pearson’s r. Statistical significance was assessed using phenotype-label permutation testing (10,000 permutations) for both analyses. Further details on CT estimation and PLSR analyses are provided in **eMethods 5**.

#### Sensitivity analysis

To assess the robustness of the findings to potential outliers, z-scores of whole-brain mean HE were calculated separately within each group. Participants with whole-brain HE values exceeding ±3 standard deviations from the group mean were excluded. This procedure identified HC outliers in PSYSCAN and one HC plus three CHR-P outliers in AMP SCZ.

Group-level statistical analyses were performed in Python v3.12.2, whereas prediction modelling was conducted in R v4.2.2.

## 3. Results

### Baseline HE differences between CHR-P individuals and HCs

#### PSYSCAN cohort

rsfMRI data were available from 190 CHR-P individuals and 115 HCs at baseline, 124 CHR-P and 69 HCs at 6 months, and 116 CHR-P and 73 HCs at 12 months. Comprehensive demographic and clinical characteristics are presented in **Table 1**.

**Table 1.**
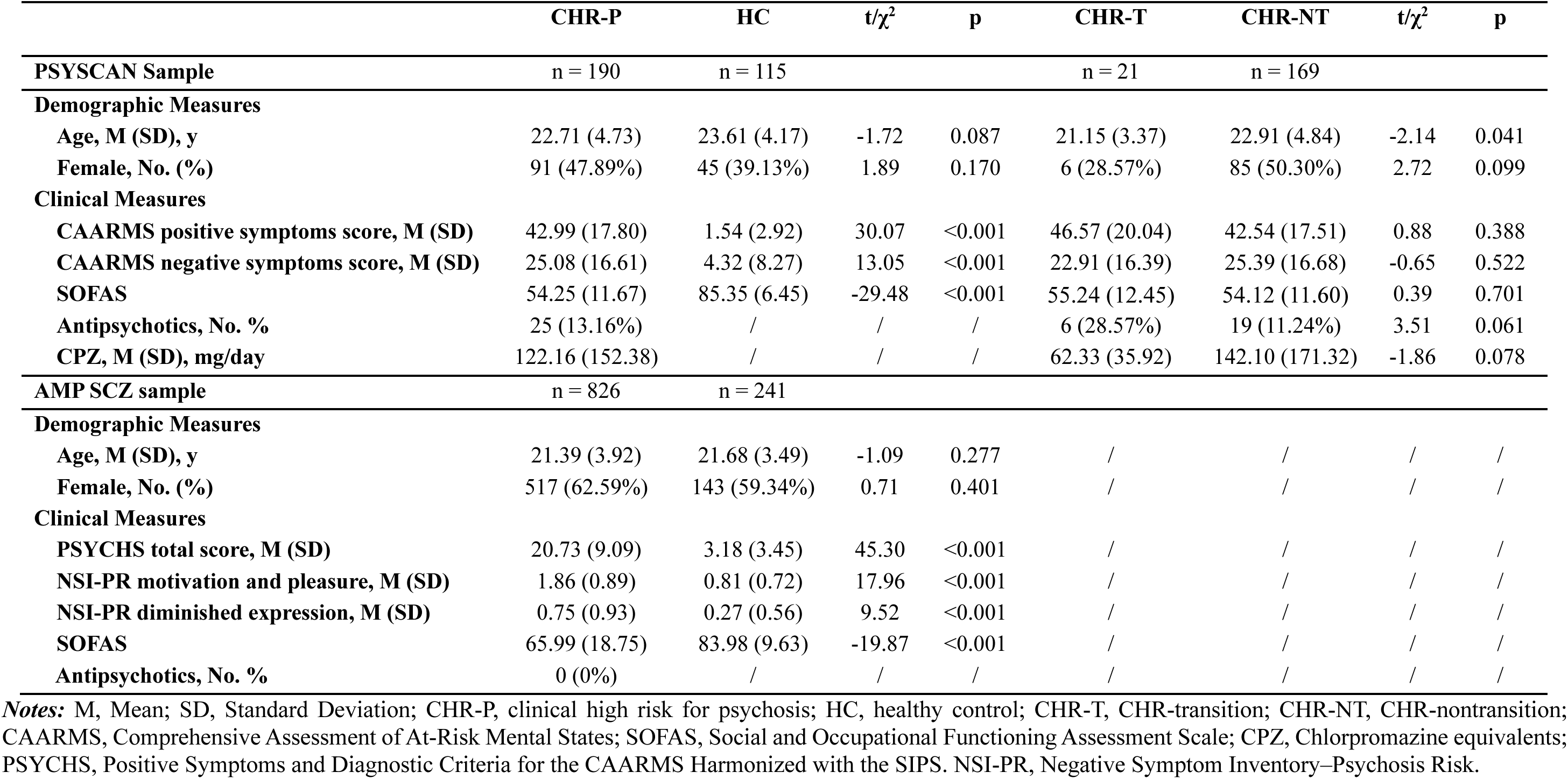
Demographic and clinical characteristics among groups in PSYSCAN and AMP SCZ.

|  | **CHR-P** | **HC** | **t/χ<sup>2</sup>** | **p** | **CHR-T** | **CHR-NT** | **t/χ<sup>2</sup>** | **p** |
|---|---|---|---|---|---|---|---|---|
| **PSYSCAN Sample** | n = 190 | n = 115 |  |  | n = 21 | n = 169 |  |  |
| **Demographic Measures** |  |  |  |  |  |  |  |  |
| **Age, M (SD), y** | 22.71 (4.73) | 23.61 (4.17) | -1.72 | 0.087 | 21.15 (3.37) | 22.91 (4.84) | -2.14 | 0.041 |
| **Female, No. (%)** | 91 (47.89%) | 45 (39.13%) | 1.89 | 0.170 | 6 (28.57%) | 85 (50.30%) | 2.72 | 0.099 |
| **Clinical Measures** |  |  |  |  |  |  |  |  |
| **CAARMS positive symptoms score, M (SD)** | 42.99 (17.80) | 1.54 (2.92) | 30.07 | <0.001 | 46.57 (20.04) | 42.54 (17.51) | 0.88 | 0.388 |
| **CAARMS negative symptoms score, M (SD)** | 25.08 (16.61) | 4.32 (8.27) | 13.05 | <0.001 | 22.91 (16.39) | 25.39 (16.68) | -0.65 | 0.522 |
| **SOFAS** | 54.25 (11.67) | 85.35 (6.45) | -29.48 | <0.001 | 55.24 (12.45) | 54.12 (11.60) | 0.39 | 0.701 |
| **Antipsychotics, No. %** | 25 (13.16%) | / | / | / | 6 (28.57%) | 19 (11.24%) | 3.51 | 0.061 |
| **CPZ, M (SD), mg/day** | 122.16 (152.38) | / | / | / | 62.33 (35.92) | 142.10 (171.32) | -1.86 | 0.078 |
| **AMP SCZ sample** | n = 826 | n = 241 |  |  |  |  |  |  |
| **Demographic Measures** |  |  |  |  |  |  |  |  |
| **Age, M (SD), y** | 21.39 (3.92) | 21.68 (3.49) | -1.09 | 0.277 | / | / | / | / |
| **Female, No. (%)** | 517 (62.59%) | 143 (59.34%) | 0.71 | 0.401 | / | / | / | / |
| **Clinical Measures** |  |  |  |  |  |  |  |  |
| **PSYCHS total score, M (SD)** | 20.73 (9.09) | 3.18 (3.45) | 45.30 | <0.001 | / | / | / | / |
| **NSI-PR motivation and pleasure, M (SD)** | 1.86 (0.89) | 0.81 (0.72) | 17.96 | <0.001 | / | / | / | / |
| **NSI-PR diminished expression, M (SD)** | 0.75 (0.93) | 0.27 (0.56) | 9.52 | <0.001 | / | / | / | / |
| **SOFAS** | 65.99 (18.75) | 83.98 (9.63) | -19.87 | <0.001 | / | / | / | / |
| **Antipsychotics, No. %** | 0 (0%) | / | / | / | / | / | / | / |

***Notes:*** M, Mean; SD, Standard Deviation; CHR-P, clinical high risk for psychosis; HC, healthy control; CHR-T, CHR-transition; CHR-NT, CHR-nontransition; CAARMS, Comprehensive Assessment of At-Risk Mental States; SOFAS, Social and Occupational Functioning Assessment Scale; CPZ, Chlorpromazine equivalents; PSYCHS, Positive Symptoms and Diagnostic Criteria for the CAARMS Harmonized with the SIPS. NSI-PR, Negative Symptom Inventory–Psychosis Risk.

At a global level, CHR-P individuals showed lower whole-brain mean HE than HCs (t = -2.09, p = 0.038, adjusted Cohen’s d = -0.250; **Figure 1**). Whole-brain HE did not differ by antipsychotic exposure status (t = −1.84, p = 0.067; **eFigure 4**).

**Figure 1.**
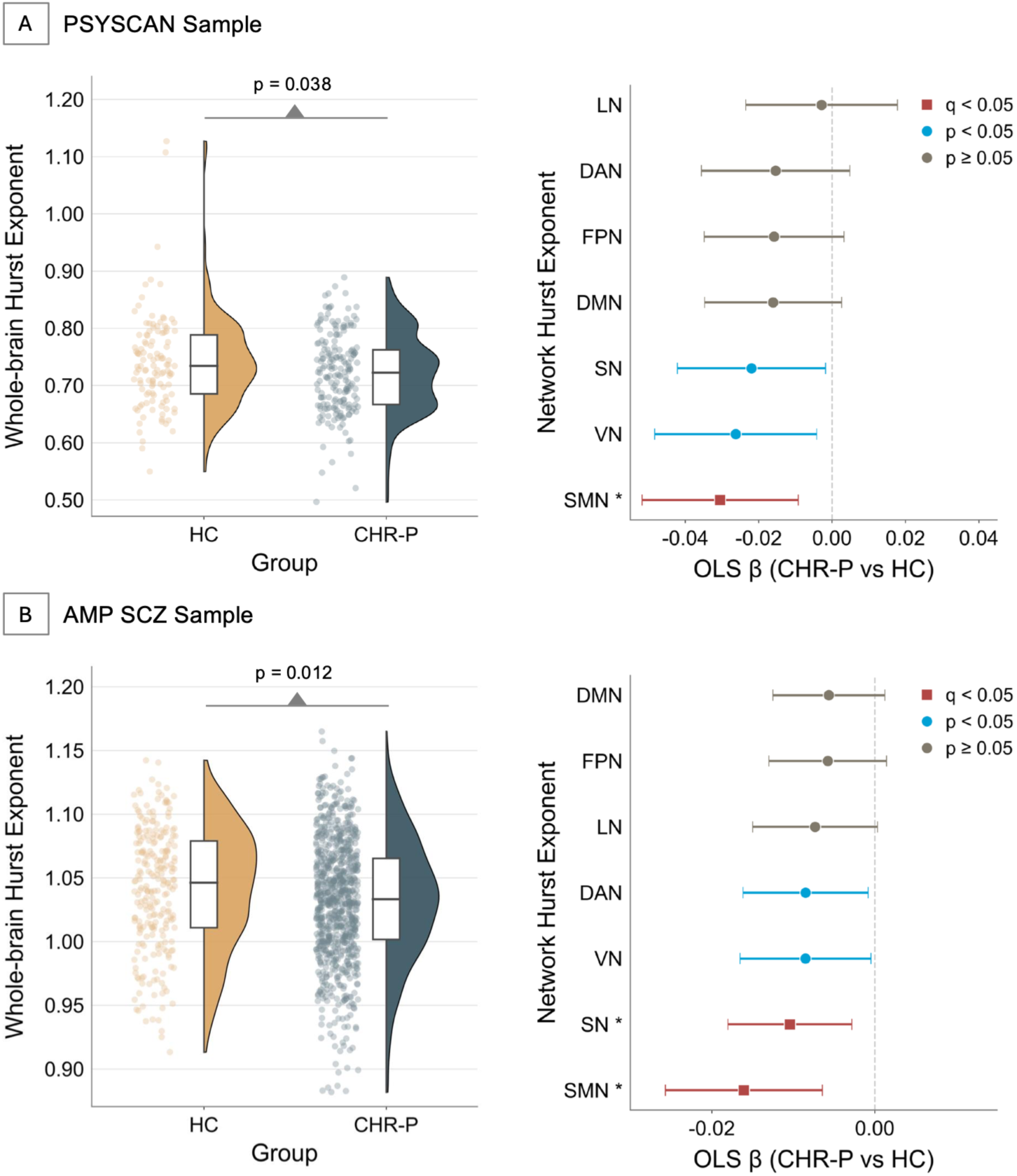
Phenotype effects on whole-brain and network-level Hurst exponent. A) Whole-brain and network-level Hurst exponent (HE) differences between clinical high risk for psychosis (CHR-P) participants and healthy controls (HCs) in the PSYSCAN sample. B) Corresponding whole-brain and network-level results in the AMP SCZ sample. In the whole-brain plots, each point represents an individual participant’s whole-brain mean HE value, adjusted for age, sex, and mean framewise displacement (MeanFD). Half-violin plots show the group-wise distributions, and box plots show the median and interquartile range. In the network-level forest plots, points represent ordinary least squares regression coefficients (OLS β) for the CHR-P versus HC contrast, with horizontal error bars indicating 95% confidence intervals. Red squares indicate effects surviving Benjamini–Hochberg false discovery rate (FDR) correction at q < 0.05, blue circles indicate nominally significant effects at p < 0.05, and gray circles indicate nonsignificant effects. Asterisks mark networks surviving FDR correction. DAN, dorsal attention network; DMN, default mode network; FPN, frontoparietal control network; LN, limbic network; SMN, somatomotor network; SN, salience/ventral attention network; VN, visual network.

At a network level, lower HE was significant in the somatomotor network (SMN, t = -2.82, p = 0.005, q = 0.035, adjusted Cohen’s d = -0.336) after FDR-correction, and in the visual network (VN, t = -2.34, p = 0.020, q = 0.070, adjusted Cohen’s d = -0.279) and the salience network (SN, t = -2.14, p = 0.033, q = 0.078, adjusted Cohen’s d = -0.255) at an uncorrected level (**Figure 1**).

At a regional level, CHR-P group had lower HE in 81 [22.5%] of the cortical parcels at an uncorrected threshold (**eTable 1**; **Figure 2),** but only one parcel (left ventral visual area V8) survived FDR-correction. Lower HE was also evident in 4 [6.1%] of the subcortical parcels (**eTable 2**; **Figure 2),** all located in the thalamus, but none of these survived FDR-correction.

**Figure 2.**
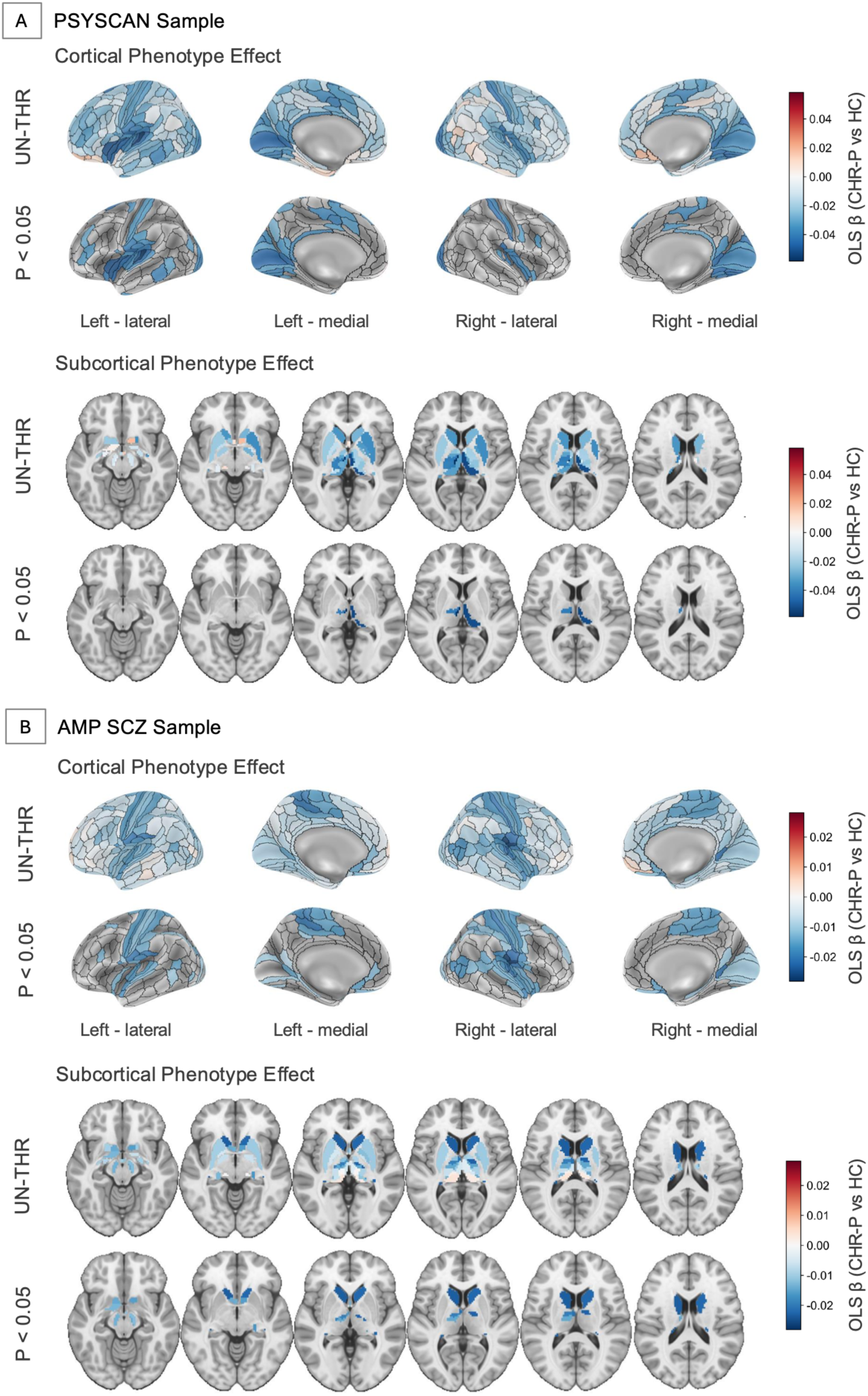
Phenotype effects on parcel-wise Hurst exponent. A) Cortical and subcortical maps showing regional Hurst exponent (HE) differences between clinical high risk for psychosis (CHR-P) participants and healthy controls (HCs) in the PSYSCAN sample. B) Corresponding cortical and subcortical maps in the AMP SCZ sample. Color bars represent ordinary least squares regression coefficients (OLS β) for the CHR-P versus HC contrast. Blue colors indicate regions with lower HE in CHR-P participants than in HCs, consistent with increased excitation. For each sample, unthresholded maps and nominally significant maps thresholded at p < 0.05 are shown.

Sensitivity analyses excluding participants with whole-brain HE values exceeding ± 3 SD from group means showed consistent directionality and patterns (**eFigure 5**). The largest network-level difference remained in the somatomotor network (SMN, t = −2.34, p = 0.020, q = 0.138; adjusted Cohen’s d = −0.281), although the whole-brain HE difference was no longer statistically significant (t = −1.46, p = 0.146; adjusted Cohen’s d = −0.180).

#### AMP SCZ cohort

rsfMRI data were available from 826 CHR-P individuals and 241 HCs at baseline, and from 385 CHR-P and 54 HCs of the baseline samples at month 2 (**Table 1**).

At a global level, CHR-P individuals showed lower whole-brain mean HE than HCs (t = -2.52, p = 0.012, adjusted Cohen’s d = -0.185; **Figure 1**).

At a network level, lower HE was significant in the SMN (t = -3.27, p = 0.001, q = 0.008, adjusted Cohen’s d = -0.241) and the SN (t = -2.69, p = 0.007, q = 0.025, adjusted Cohen’s d = -0.198) after FDR-correction, and in the VN (t = -2.05, p = 0.041, q = 0.067, adjusted Cohen’s d = -0.153) and dorsal attention network (DAN, t = -2.17, p = 0.030, q = 0.067, adjusted Cohen’s d = -0.160; **Figure 1**) at an uncorrected level.

At a regional level, CHR-P individuals showed lower HE in 120 cortical parcels [33.3%] at an uncorrected threshold, and in 20 cortical parcels [5.6%] after FDR-correction (**eTable 1**; **Figure 2).** The latter were located in the posterior opercular, auditory, somatosensory-motor, posterior cingulate, paracentral lobular/mid-cingulate, superior parietal, and anterior cingulate/medial prefrontal cortices. Lower HE was also observed in 23 [34.8%] of the subcortical parcels, 9 of which [13.6%] survived FDR-correction (**eTable 2**; **Figure 2).** These were mainly in the thalamus, caudate, and ventral tegmental area (VTA).

Sensitivity analysis indicated that after exclusion of outliers, the group effects became larger for both whole-brain (t = -2.61, p = 0.009, adjusted Cohen’s d = -0.193) and network-level HE, with the most prominent effects in the SMN (t = -3.36, p = 0.001, q = 0.006, adjusted Cohen’s d = -0.248), salience network (SN, t = -2.78, p = 0.006, q = 0.020, adjusted Cohen’s d = -0.205), visual network (VN, t = -2.21, p = 0.0276, q = 0.048, adjusted Cohen’s d = -0.163) and dorsal attention network (DAN, t = -2.30, p = 0.022, q = 0.048, adjusted Cohen’s d = -0.170; **eFigure 5**).

Across both cohorts, the most prominent reductions in HE were observed in the SMN, with additional reductions in the SN and VN. At the subcortical level, the largest reductions were consistently observed in the thalamus. We further examined the spatial correlation of regional case–control effects between the two cohorts. The cortical effect maps showed significant spatial correspondence (r = 0.26, p_permuted =_ 0.032; **eFigure 6**), and the subcortical maps did not (r = -0.18, p_permuted =_ 0.346).

### Longitudinal HE differences between CHR-P individuals and HCs

#### PSYSCAN cohort

At a global level, HE did not show a significant phenotype-by-time interaction (F (2, 440.09) = 1.85, p = 0.159; **Figure 3**). Similarly, no significant longitudinal group differences were identified at the network level (p > 0.05; **eFigure 7**), and none of the parcel-level effects survived FDR-correction (q > 0.05).

**Figure 3.**
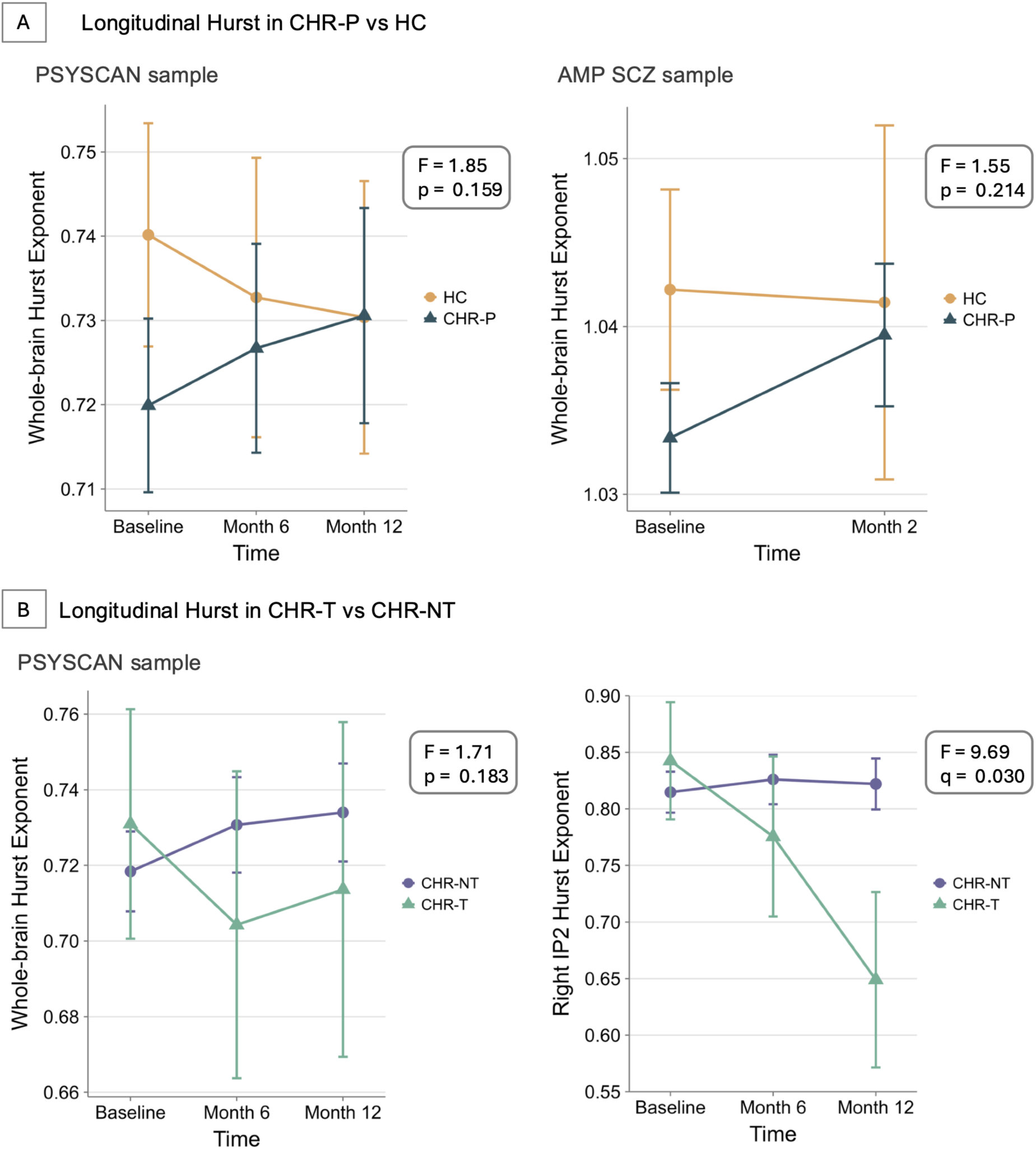
Group differences in longitudinal trajectories of Hurst exponent. A) Longitudinal trajectories of whole-brain Hurst exponent (HE) in clinical high risk for psychosis (CHR-P) participants and healthy controls (HCs) in the PSYSCAN and AMP SCZ samples. B) Longitudinal trajectories of whole-brain HE and right inferior parietal IP2 parcel in transition (CHR-T) and non-transition (CHR-NT) participants in the PSYSCAN sample. Points represent model-estimated marginal means adjusted for age, sex, and mean framewise displacement (MeanFD), and vertical error bars indicate 95% confidence intervals. Lines connect estimated marginal means across time points. Model statistics indicate the group-by-time interaction effect from linear mixed-effects models. Baseline, Month 6, and Month 12 were available in the PSYSCAN sample, while baseline and Month 2 were available in the AMP SCZ sample.

#### AMP SCZ cohort

No significant phenotype-by-time interactions were observed at the whole-brain (F (1, 598.38) = 1.55, p = 0.214; **Figure 3**), network (p > 0.05; **eFigure 7**), or parcel levels (q > 0.05).

#### Baseline HE differences between CHR-T and CHR-NT individuals

In PSYSCAN, twenty-one CHR-P participants (11%) developed psychosis during follow-up (mean follow-up = 724.1 days; standard deviation (SD) = 443.7 days). From this CHR-T subgroup, rsfMRI data were available from 21 subjects at baseline, 11 subjects at 6 months, and 9 subjects at 12 months (**Table 1**).

At a global level, HE did not differ between CHR-T and CHR-NT individuals (t = 0.44, p = 0.660; **eFigure 8**). At a network level, no significant group differences were observed (all p > 0.05). At a regional level, nominal differences in HE were observed in 11 cortical parcels [3.1%] and 2 subcortical parcels [3.0%], although none survived FDR-correction (**eFigure 8**; **eTable 3**). These parcels included the anterior cingulate/medial prefrontal, early auditory, medial temporal, paracentral lobule/mid-cingulate, dorsolateral prefrontal, superior parietal cortices, and the substantia nigra and VTA.

We also assessed the ability of baseline regional HE to predict transition to psychosis at the individual level. The elastic-net Cox model showed above-chance discrimination, with a Harrell’s C-index of 0.63 (95% CI : 0.59–0.68). It achieved a BAC of 0.62 (95% CI : 0.58–0.67), with sensitivity of 0.83 (95% CI : 0.74–0.93), specificity of 0.41 (95% CI : 0.26–0.57) and Brier score of 0.11 (95% CI : 0.10-0.13). Most predictors were shrunk to zero by the elastic-net penalty, while the retained regional HE features showed notable overlap with regions exhibiting nominally significant differences between CHR-T and CHR-NT (**eTable 4**).

#### Longitudinal HE differences between CHR-T and CHR-NT individuals

At a whole-brain level, the longitudinal trajectories of HE did not differ significantly between the CHR-T and CHR-NT groups (F (2, 279.30) = 1.71, p = 0.183; **Figure 3**). There was also an absence of significant longitudinal differences at a network level (p > 0.05; **eFigure 9**). However, visual inspection indicated that, at both global and network levels, HE progressively decreased in the CHR-T group but remained stable or increased in the CHR-NT group.

At the regional level, a significant phenotype-by-time interaction was identified in the right inferior parietal IP2 parcel (F (2, 294.52) = 9.69, p < 0.001, q = 0.030; **Figure 3**); no other parcels survived correction. In this region, HE was lower at both follow-up time points than at baseline in the CHR-T group, but remained relatively stable over time in the CHR-NT group.

#### Biological substrates of HE phenotype effect

Having identified significant cross-cohort correspondence of cortical case-control effects on HE, we also examined their associations with structural and molecular features.

Regional case-control effects on HE were not spatially associated with CT effects in PSYSCAN after accounting for spatial autocorrelation using permutation testing (r = -0.03, p_permuted =_ 0.716), but showed a significant negative spatial association in AMP SCZ (r = -0.18, p_permuted =_ 0.047; **eFigure 10**).

Receptor/transporter distribution patterns significantly predicted regional HE effects in both the PSYSCAN (r = 0.52, p_permuted =_ 0.048; **Figure 4**) and AMP SCZ (r = 0.62, p_permuted =_ 0.036). The strongest contributors (variable importance in projection [VIP]>1) were opioid (MOR), norepinephrine (NET), serotonin (5-HT_1A,_ 5-HTT, 5-HT₄), cannabinoid (CB_1)_, and GABAa in PSYSCAN, and norepinephrine (NET), serotonin (5-HTT, 5-HT₂_A_, 5-HT₄), acetylcholine (VAChT), dopamine (DAT), and opioid (MOR) in AMP SCZ.

**Figure 4.**
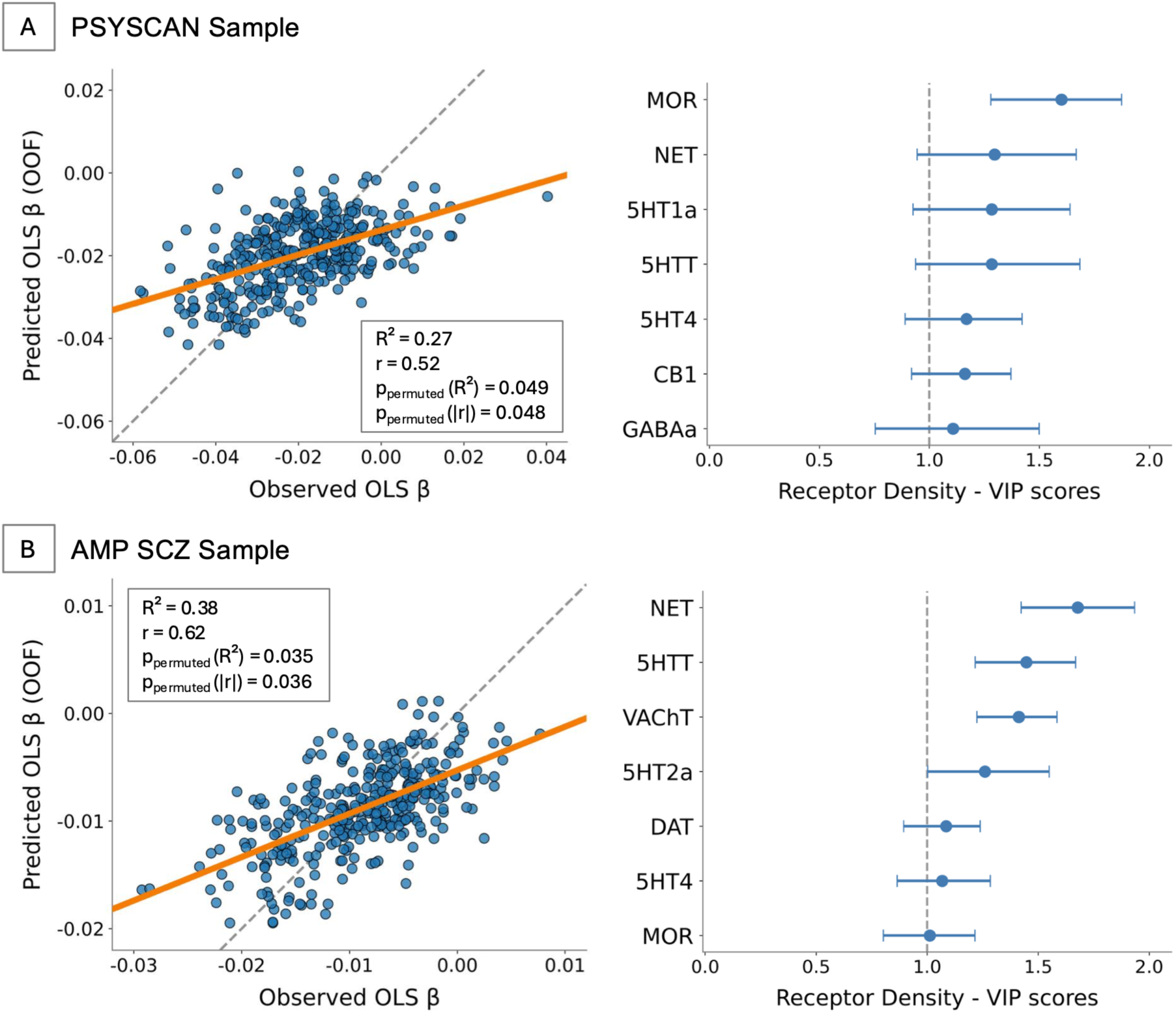
Receptor-density prediction of phenotype-associated Hurst exponent effects. Results from A) the PSYSCAN sample and B) the AMP SCZ sample. For each sample, scatter plots show predicted versus observed parcel-wise phenotype-associated Hurst exponent ordinary least squares regression coefficients (OLS β) from 10-fold cross-validated partial least squares regression (PLSR) based on receptor density. Each point represents one cortical parcel. The solid orange line shows the best-fit regression line, and the dashed gray line indicates the identity line (y = x), representing perfect agreement between predicted and observed values. Model performance is summarized using out-of-fold prediction accuracy, including R², Pearson’s r, and permutation-based p-values for R² and |r|. Forest plots show variable importance in projection (VIP) scores from the PLSR model. Points represent bootstrap mean VIP scores, and horizontal error bars indicate 95% confidence intervals. The dashed vertical line marks VIP = 1. Receptors with VIP > 1 were considered to contribute above average to the model. MOR, μ-opioid receptor; NET, norepinephrine transporter; 5HT1A, 5HT4, and 5HT2A, serotonin receptors; 5HTT, serotonin transporter; CB1, cannabinoid receptor type 1; GABAA, gamma-aminobutyric acid type A receptor; VAChT, vesicular acetylcholine transporter; DAT, dopamine transporter.

## 4. Discussion

To our knowledge, this is the first investigation of E/I balance in CHR-P individuals using fMRI. We found lower HE in CHR-P individuals than controls in one cohort, then replicated this finding in a second independent cohort. Within the CHR-P group, longitudinal analyses provided preliminary evidence of a transition-related reduction in HE, and baseline regional HE patterns also carried modest individual-level prognostic information for subsequent transition. Collectively, these results suggest that increased vulnerability to psychosis is associated with reduced HE, an index of E/I imbalance.

Both CHR-P cohorts showed altered HE in a similar topographic pattern, primarily involving somatomotor, salience, and visual cortical networks and thalamic nuclei. Lower HE suggests a net increase in excitation, consistent with prior magnetoencephalographic (MEG) and MRS data in CHR-P participants.^36–38^ The distribution of HE alterations aligns with sensory gating and integration deficits,^39,40^ aberrant salience processing,^41^ somato-visual dysconnectivity,^42^ and disrupted thalamocortical circuits in psychosis.^43,44^ The involvement of somatomotor cortex and the thalamus is also consistent with preclinical evidence that PV interneurons are enriched in sensory/motor cortices and participate in feed-forward inhibitory circuits gating information flow within thalamocortical systems.^45–47^ It’s also in line with the topographic distribution of lower HE in patients with first-episode psychosis.^17^

Within the CHR-P group, we observed only nominal regional HE differences at the group level related to transition to psychosis, although individual-level models based on baseline regional HE showed above-chance predictive performance. This discrepancy may reflect heterogeneity in the mechanisms underlying psychosis onset,^48^ which could make effects harder to detect when analysing mean data from a group. A further consideration is that multivariable models can integrate complementary information across regions not captured by separate region-wise comparisons. The regions showing the largest transition-related group differences and contributing to prediction included the prefrontal/anterior cingulate cortices, substantia nigra and VTA. These areas are of particular interest given previous evidence linking subcortical dopamine dysfunction and alterations in prefrontal and midbrain activation and connectivity to psychosis onset in CHR-P samples.^49–51^

The spatial distribution of HE alterations did not positively align with CT differences, suggesting that HE doesn’t simply reflect consistent macroscopic structural variation. However, in both cohorts, HE alterations were linked to the normative distribution of norepinephrine (NET), serotonin (5-HT), and opioid (MOR) receptors. Norepinephrine has dose-dependent modulatory effects on sensory processing in somatosensory cortex, partly through its influence on E/I balance,^52^ and has been implicated in cognitive impairments in psychosis.^53^ Mouse-model studies suggest that dysregulated 5-HT₂A receptor signaling may contribute to cortical hyperexcitability in psychosis by impairing PV interneuron–mediated GABAergic inhibition.^54^ MOR availability has also been found to be reduced in schizophrenia, and its activation can inhibit GABA release, potentially reducing inhibition.^55,56^

Beyond these mechanistic insights, HE offers translational potential for early detection of psychosis risk. As a rsfMRI-derived measure of intrinsic neural dynamics, HE is task-free, non-invasive, and suitable for the assessment of large participant samples across multiple sites.^12^ By indexing E/I-related alterations, HE may support neurobiologically informed CHR-P phenotyping and capture macroscale, system-level brain vulnerability associated with psychosis risk. Its potential applicability across species^14^ may help bridge human neuroimaging findings with preclinical models of E/I dysfunction, supporting mechanistically informed biomarker development for precision psychiatry.

## Limitations

HE is an indirect proxy of E/I balance, and multimodal studies combining HE with neurochemical or metabolic measures, such as PET or MRS, or with pharmacological manipulations may further validate this link. Methodological studies are also needed to establish the stability of HE across acquisition and processing choices. In PSYSCAN, the small number of CHR-P participants who developed psychosis limited our power to detect HE correlates of transition. Finally, effect sizes for case-control differences were modest, consistent with evidence that neuroimaging and cognitive alterations at the CHR-P stage are less pronounced than in psychotic patients,^57,58^ and with our recent finding of larger HE alterations in first-episode psychosis.^17^

## Conclusion

In two large cohorts of people at CHR-P, we found evidence of distributed changes in HE that may indicate a shift towards excitation relative to inhibition, particularly in somatomotor and salience networks, and in the thalamus. The distribution of these alterations was related to neurotransmitter receptor distributions, particularly those of norepinephrine, serotonin, and opioid systems. The pattern of alterations in HE at the individual level may be helpful for predicting the later onset of psychosis.

## Data availability

Data from the PSYSCAN study are available through an approved project proposal submitted to the PSYSCAN Board (contact: Dr Alexis Cullen). The AMP SCZ dataset used in this study was accessed through the National Institute of Mental Health Data Archive (NDA; Release 3.0; DOI: 10.15154/81xe-k706) under an approved Data Use Certification. AMP SCZ data are available via scheduled releases at the NDA AMP SCZ Data Repository (https://nda.nih.gov/ampscz).

## Acknowledgement

The PSYSCAN project is supported by grant agreement No. 603196 under the European Union’s Seventh Framework Programme.

This paper uses data collected in the Accelerating Medicines Partnership® Schizophrenia (AMP® SCZ) project. AMP SCZ is supported by NIMH grants (U24MH124629, U01MH124631, U01MH124639) and by the Wellcome Trust (220664/Z/20/Z).

Li and Oliver are funded by the Wellcome Trust (STEP, 216005/Z/19/Z). Varvari, Oliver, McCutcheon, McGuire are supported by the NIHR Oxford Health Biomedical Research Centre. The views expressed are those of the author(s) and not necessarily those of the NHS, the NIHR or the Department of Health.

## PSYSCAN Study Group Authors

Matthew J. Kempton¹, Kate Merritt¹^˒^², Andrea Mechelli¹, Natalia Petros¹, Mathilde Antoniades¹, Andrea De Micheli¹, Sandra Vieira¹^˒^³^˒^⁴, Tom Spencer¹^˒^⁵, Frederike Schirmbeck⁶, Diana Tordesillas-Gutierrez⁷^˒^⁸, Esther Setien-Suero⁷^˒^⁸, Rosa Ayesa-Arriola⁷^˒^⁸, Paula Suarez-Pinilla⁷^˒^⁸, Victor Ortiz Garcia-de la foz⁷^˒^⁸, Mikkel Erlang Sørensen⁹, Bjørn H. Ebdrup⁹^˒^¹⁰, Karen Tangmose⁹^˒^¹⁰, Helle Schæbel⁹, Egill Rostrup⁹^˒^¹¹, Anja Richter¹², Bernd Krämer¹², Bea Campforts¹³, Machteld Marcelis¹³^˒^¹⁴, Claudia Vingerhoets¹³, Covadonga Díaz-Caneja¹⁵, Miriam Ayora¹⁵, Joost Janssen¹⁶, Roberto Rodriguez-Jimenez¹⁷, Marina Díaz-Marsá¹⁸, Irina Falkenberg¹⁹, Florian Bitsch¹⁹, Jens Sommer¹⁹, Patrick McGorry^²⁰˒²¹^, Paul Amminger²⁰^˒^²¹, Meredith McHugh^²⁰˒²¹^, Armida Mucci²², Paola Bucci²², Giuseppe Piegari²², Daria Pietrafesa²², Sara Patriarca²², André Zugman²³, Ary Gadelha²³, Graccielle Rodrigues da Cunha²³, Kang Ik Kevin Cho²⁴^˒^²⁵, Taeyoung Lee²⁶^˒^²⁷, Minah Kim²⁸^˒^²⁹, Sun Young Moon²⁸^˒^³⁰, Silvia Kyungjin Lho²⁸^˒^³¹, Michael Kiang³²^˒^³³, Cory Gerritsen³³^˒^³⁴, Margaret Maheandiran³³, Sarah Ahmed³⁵^˒^³³, Jenny Lepock³⁵^˒^³³, Matthäus Willeit³⁶, Marzena Lenczowska³⁶, Ulrich Sauerzopf³⁶, Ana Weidenauer³⁶, Julia Furtner³⁷, Achim Burrer³⁸, Philipp Stämpfli³⁸, Naemi Huber³⁸, Wolfram Kawohl³⁹.

## Affiliations of PSYSCAN Study Group Authors

**1.** Department of Psychosis Studies, Institute of Psychiatry, Psychology & Neuroscience, King’s College London, De Crespigny Park, Denmark Hill, London, United Kingdom SE5 8AF.

**2.** MRC Centre for Neurodevelopmental Disorders, King’s College London, UK.

**3.** Department of Radiology, Lausanne University Hospital and University of Lausanne (CHUV-UNIL), Lausanne, Switzerland.

**4.** Center for Research in Neuropsychology and Cognitive Behavioral Intervention, Faculty of Psychology and Educational Sciences, University of Coimbra, Coimbra, Portugal.

**5.** Outreach and Support in South-London (OASIS) service, South London and Maudsley (SLaM) NHS Foundation Trust, UK.

**6.** Amsterdam UMC, University of Amsterdam, Psychiatry, Department Early Psychosis, Meibergdreef 9, Amsterdam, The Netherlands.

**7.** Department of Psychiatry, Marqués de Valdecilla University Hospital, IDIVAL. School of Medicine, University of Cantabria, Santander, Spain.

**8.** CIBERSAM, Centro Investigación Biomédica en Red Salud Mental, Spain.

**9.** Centre for Neuropsychiatric Schizophrenia Research (CNSR) & Centre for Clinical Intervention and Neuropsychiatric Schizophrenia Research (CINS), Mental Health Centre Glostrup, University of Copenhagen, Glostrup, Denmark.

**10.** University of Copenhagen, Faculty of Health and Medical Sciences, Dept. of Clinical Medicine, Copenhagen, Denmark.

**11.** Functional Imaging Unit (FIUNIT), Rigshospitalet Glostrup, University of Copenhagen, Glostrup, Denmark.

**12.** Section for Experimental Psychopathology and Neuroimaging, Department of General Psychiatry, Heidelberg University, Heidelberg, Germany.

**13.** Department of Psychiatry and Neuropsychology, Maastricht University, Maastricht, The Netherlands.

**14.** GGZE Mental Health Care, Eindhoven, the Netherlands.

**15.** Department of Child and Adolescent Psychiatry, Institute of Psychiatry and Mental Health, Hospital General Universitario Gregorio Marañón, IiSGM, CIBERSAM, ISCIII, School of Medicine, Universidad Complutense, Madrid, Spain.

**16.** Department of Child and Adolescent Psychiatry, Institute of Psychiatry and Mental Health, Hospital General Universitario Gregorio Marañón, IiSGM, CIBERSAM, ISCIII, Madrid, Spain.

**17.** Department of Psychiatry, Instituto de Investigación Sanitaria Hospital 12 de Octubre (imas 12), CIBERSAM-ISCIII, School of Medicine, Universidad Complutense (UCM), Madrid, Spain.

**18.** Department of Psychiatry, Instituto de Investigación Hospital Clínico San Carlos (IdISSC), CIBERSAM, ISCIII, School of Medicine, Universidad Complutense, Madrid, Spain.

**19.** Department of Psychiatry, University of Marburg, Rudolf-Bultmann-Straße 8, D-35039, Marburg, Germany.

**20.** Orygen, 35 Poplar Road, Parkville, Victoria, Melbourne, Australia.

**21.** Centre for Youth Mental Health, The University of Melbourne, Parkville, Victoria, Australia.

**22.** Department of Psychiatry, University of Campania Luigi Vanvitelli, Largo Madonna delle Grazie, 80138, Naples, Italy.

**23.** Department of Psychiatry, Interdisciplinary Lab for Clinical Neurosciences (LiNC), Universidade Federal de Sao Paulo (UNIFESP), Sao Paulo, Brazil.

**24.** Department of Brain and Cognitive Sciences, Seoul National University College of Natural Sciences, Gwanakro1, Gwanak-gu, Seoul, Republic of Korea.

**25.** Department of Psychiatry, Brigham and Women’s Hospital, Harvard Medical School, Boston, MA, USA.

**26.** Department of Psychiatry, Kyungpook National University School of Medicine, Daegu, Republic of Korea.

**27.** Institute of Human Behavioral Medicine, SNU-MRC, 101 Daehakno, Jongno-gu, Seoul, Republic of Korea.

**28.** Department of Psychiatry, Seoul National University College of Medicine, 101 Daehakno, Jongno-gu, Seoul, Republic of Korea.

**29.** Department of Neuropsychiatry, Seoul National University Hospital, 101 Daehakno, Jongno-gu, Seoul, Korea.

**30.** Department of Public Health Medical Services, Seoul National University Bundang Hospital, Seongnam, Republic of Korea.

**31.** Department of Psychiatry, Seoul Metropolitan Government-Seoul National University Boramae Medical Center, Seoul, Republic of Korea.

**32.** Department of Psychiatry, University of Toronto, 250 College St 8th Floor, Toronto, Ontario, Canada M5T 1R8.

**33.** Centre for Addiction and Mental Health, 250 College Street, Toronto, Ontario, Canada M5T 1R8.

**34.** Department of Psychology, University of Toronto, 100 St. George Street 4th Floor, Toronto, Ontario, Canada M5S 3G3.

**35.** Institute of Medical Science, University of Toronto, 1 King’s College Circle Room 2374, Toronto, Ontario, Canada M5S 1A8.

**36.** Department of Psychiatry and Psychotherapy, 1090 Vienna, Austria.

**37.** Medical University of Vienna, Department of Biomedical Imaging and Image-guided Therapy Währingergürtel 18-20, 1090 Vienna.

**38.** Department of Psychiatry, Psychotherapy and Psychosomatics, Psychiatric Hospital, University of Zurich, Switzerland.

**39.** Department for Psychiatry and Psychotherapy, Psychiatric Services Aargau, Brugg, Switzerland.

## References

1. Stahl SM. Beyond the dopamine hypothesis of schizophrenia to three neural networks of psychosis: dopamine, serotonin, and glutamate. CNS Spectr. 2018;23(3):187–191. doi:10.1017/S1092852918001013

2. Moghaddam B, Javitt D. From revolution to evolution: the glutamate hypothesis of schizophrenia and its implication for treatment. Neuropsychopharmacology. 2012;37(1):4–15. doi:10.1038/npp.2011.181

3. Lewis DA, Curley AA, Glausier JR, Volk DW. Cortical parvalbumin interneurons and cognitive dysfunction in schizophrenia. Trends Neurosci. 2012;35(1):57–67. doi:10.1016/j.tins.2011.10.004

4. Jami SA, Cameron S, Wong JM, Daly ER, McAllister AK, Gray JA. Increased excitation-inhibition balance and loss of GABAergic synapses in the serine racemase knockout model of NMDA receptor hypofunction. J Neurophysiol. 2021;126(1):11–27. doi:10.1152/jn.00661.2020

5. Pafundo DE, Pretell Annan CA, Fulginiti NM, Belforte JE. Early NMDA receptor ablation in interneurons causes an activity-dependent E/I imbalance in vivo in prefrontal cortex pyramidal neurons of a mouse model useful for the study of schizophrenia. Schizophr Bull. 2021;47(5):1300–1309. doi:10.1093/schbul/sbab030

6. Isaacson JS, Scanziani M. How inhibition shapes cortical activity. Neuron. 2011;72(2):231–243. doi:10.1016/j.neuron.2011.09.027

7. Anticevic A, Lisman J. How can global alteration of excitation/inhibition balance lead to the local dysfunctions that underlie schizophrenia? Biol Psychiatry. 2017;81(10):818–820. doi:10.1016/j.biopsych.2016.12.006

8. Howes O, McCutcheon R, Stone J. Glutamate and dopamine in schizophrenia: an update for the 21st century. J Psychopharmacol. 2015;29(2):97–115. doi:10.1177/0269881114563634

9. Rawani NS, Chan AW, Dursun SM, Baker GB. The underlying neurobiological mechanisms of psychosis: focus on neurotransmission dysregulation, neuroinflammation, oxidative stress, and mitochondrial dysfunction. Antioxidants (Basel). 2024;13(6):709. doi:10.3390/antiox13060709

10. Jardri R, Hugdahl K, Hughes M, et al. Are hallucinations due to an imbalance between excitatory and inhibitory influences on the brain? Schizophr Bull. 2016;42(5):1124–1134. doi:10.1093/schbul/sbw075

11. Campbell OL, Weber AM. Monofractal analysis of functional magnetic resonance imaging: An introductory review. Hum Brain Mapp. 2022;43(8):2693–2706. doi:10.1002/hbm.25801

12. Trakoshis S, Martínez-Cañada P, Rocchi F, et al. Intrinsic excitation-inhibition imbalance affects medial prefrontal cortex differently in autistic men versus women. eLife. 2020;9:e55684. doi:10.7554/eLife.55684

13. Xie K, Royer J, Rodriguez-Cruces R, et al. Temporal Lobe Epilepsy Perturbs the Brain- Wide Excitation-Inhibition Balance: Associations with Microcircuit Organization, Clinical Parameters, and Cognitive Dysfunction. Advanced Science. 2025;12(9):2406835. doi:10.1002/advs.202406835

14. Nishio M, Ellwood-Lowe ME, Woodburn M, et al. The development of neural inhibition across species: insights from the hurst exponent. J Neurosci. 2025;45(41):e0025252025. doi:10.1523/JNEUROSCI.0025-25.2025

15. Steullet P, Cabungcal JH, Coyle J, et al. Oxidative stress-driven parvalbumin interneuron impairment as a common mechanism in models of schizophrenia. Mol Psychiatry. 2017;22(7):936–943. doi:10.1038/mp.2017.47

16. Uliana DL, Lisboa JRF, Gomes FV, Grace AA. The excitatory-inhibitory balance as a target for the development of novel drugs to treat schizophrenia. Biochemical Pharmacology. 2024;228:116298. doi:10.1016/j.bcp.2024.116298

17. Varvari I, Doody M, Li Z, et al. Convergent multimodal evidence of cortical excitation-inhibition imbalance in psychosis. bioRxiv. Preprint posted online April 6, 2026. doi:10.64898/2026.03.31.715583

18. Abbasi S, Wolff A, Çatal Y, Northoff G. Increased noise relates to abnormal excitation-inhibition balance in schizophrenia: a combined empirical and computational study. Cereb Cortex. 2023;33(20):10477–10491. doi:10.1093/cercor/bhad297

19. Uscătescu LC, Hyatt CJ, Dunn J, et al. Using the excitation/inhibition ratio to optimize the classification of autism and schizophrenia. Transl Psychiatry. 2025;15(1):234. doi:10.1038/s41398-025-03455-8

20. Salazar De Pablo G, Radua J, Pereira J, et al. Probability of transition to psychosis in individuals at clinical high risk: an updated meta-analysis. JAMA Psychiatry. 2021;78(9):970–978. doi:10.1001/jamapsychiatry.2021.0830

21. Tognin S, Van Hell HH, Merritt K, et al. Towards precision medicine in psychosis: benefits and challenges of multimodal multicenter studies—PSYSCAN: translating neuroimaging findings from research into clinical practice. Schizophr Bull. 2020;46(2):432–441. doi:10.1093/schbul/sbz067

22. Tognin S, Vieira S, Oliver D, et al. PSYSCAN multi-centre study: baseline characteristics and clinical outcomes of the clinical high risk for psychosis sample. Schizophrenia (Heidelb). 2025;11(1):66. doi:10.1038/s41537-025-00598-x

23. Wannan CMJ, Nelson B, Addington J, et al. Accelerating medicines partnership® schizophrenia (AMP® SCZ): rationale and study design of the largest global prospective cohort study of clinical high risk for psychosis. Schizophr Bull. 2024;50(3):496–512. doi:10.1093/schbul/sbae011

24. Yung AR, Yuen HP, McGorry PD, et al. Mapping the onset of psychosis: the Comprehensive Assessment of At-Risk Mental States. Aust N Z J Psychiatry. 2005;39(11-12):964–971. doi:10.1080/j.1440-1614.2005.01714.x

25. Schultze-Lutter F, Ruhrmann S, Picker H, Klosterkötter J. Development and evaluation of the schizophrenia proneness instrument, adult version (SPI-A). Schizophr Res. 2006;86:S4–S5. doi:10.1016/S0920-9964(06)70014-7

26. Woods SW, Parker S, Kerr MJ, et al. Development of the PSYCHS: Positive SYmptoms and Diagnostic Criteria for the CAARMS Harmonized with the SIPS. Early Interv Psychiatry. 2024;18(4):255–272. doi:10.1111/eip.13457

27. Goldman HH, Skodol AE, Lave TR. Revising axis V for DSM-IV: a review of measures of social functioning. Am J Psychiatry. 1992;149(9):1148–1156. doi:10.1176/ajp.149.9.1148

28. Strauss GP, Walker EF, Carter NT, Luther L, Mittal VA. The negative symptom inventory-psychosis risk (NSI-PR): psychometric validation of the final 11-item version. Schizophr Bull. 2025;51(5):1380–1389. doi:10.1093/schbul/sbae206

29. Esteban O, Markiewicz CJ, Blair RW, et al. fMRIPrep: a robust preprocessing pipeline for functional MRI. Nat Methods. 2019;16(1):111–116. doi:10.1038/s41592-018-0235-4

30. Patel AX, Kundu P, Rubinov M, et al. A wavelet method for modeling and despiking motion artifacts from resting-state fMRI time series. Neuroimage. 2014;95:287–304. doi:10.1016/j.neuroimage.2014.03.012

31. Huang CC, Rolls ET, Feng J, Lin CP. An extended Human Connectome Project multimodal parcellation atlas of the human cortex and subcortical areas. Brain Struct Funct. 2022;227(3):763–778. doi:10.1007/s00429-021-02421-6

32. Fortin JP, Cullen N, Sheline YI, et al. Harmonization of cortical thickness measurements across scanners and sites. Neuroimage. 2018;167:104–120. doi:10.1016/j.neuroimage.2017.11.024

33. Yeo BTT, Krienen FM, Sepulcre J, et al. The organization of the human cerebral cortex estimated by intrinsic functional connectivity. J Neurophysiol. 2011;106(3):1125–1165. doi:10.1152/jn.00338.2011

34. Fischl B, Dale AM. Measuring the thickness of the human cerebral cortex from magnetic resonance images. Proc Natl Acad Sci U S A. 2000;97(20):11050–11055. doi:10.1073/pnas.200033797

35. Hansen JY, Shafiei G, Markello RD, et al. Mapping neurotransmitter systems to the structural and functional organization of the human neocortex. Nat Neurosci. 2022;25(11):1569–1581. doi:10.1038/s41593-022-01186-3

36. Grent-’t-Jong T, Gross J, Goense J, et al. Resting-state gamma-band power alterations in schizophrenia reveal E/I-balance abnormalities across illness-stages. eLife. 2018;7:e37799. doi:10.7554/eLife.37799

37. Arazi A, Toso A, Grent-‘t-Jong T, Uhlhaas PJ, Donner TH. Large-scale maps of altered cortical dynamics in early-stage psychosis are related to GABAergic and glutamatergic neurotransmission. Sci Adv. 2025;11(33):eads0400. doi:10.1126/sciadv.ads0400

38. Merritt K, Egerton A, Kempton MJ, Taylor MJ, McGuire PK. Nature of glutamate alterations in schizophrenia: a meta-analysis of proton magnetic resonance spectroscopy studies. JAMA Psychiatry. 2016;73(7):665–674. doi:10.1001/jamapsychiatry.2016.0442

39. San-Martin R, Castro LA, Menezes PR, Fraga FJ, Simões PW, Salum C. Meta-analysis of sensorimotor gating deficits in patients with schizophrenia evaluated by prepulse inhibition test. Schizophr Bull. 2020;46(6):1482–1497. doi:10.1093/schbul/sbaa059

40. Carter O, Bennett D, Nash T, et al. Sensory integration deficits support a dimensional view of psychosis and are not limited to schizophrenia. Transl Psychiatry. 2017;7(5):e1118. doi:10.1038/tp.2017.69

41. Kapur S. Psychosis as a state of aberrant salience: a framework linking biology, phenomenology, and pharmacology in schizophrenia. Am J Psychiatry. 2003;160(1):13–23. doi:10.1176/appi.ajp.160.1.13

42. Keane BP, Abrham YT, Cole MW, Johnson BA, Hu B, Cocuzza CV. Functional dysconnectivity of visual and somatomotor networks yields a simple and robust biomarker for psychosis. Mol Psychiatry. 2025;30(4):1539–1547. doi:10.1038/s41380-024-02767-3

43. Murray JD, Anticevic A. Toward understanding thalamocortical dysfunction in schizophrenia through computational models of neural circuit dynamics. Schizophr Res. 2017;180:70–77. doi:10.1016/j.schres.2016.10.021

44. Woodward ND, Karbasforoushan H, Heckers S. Thalamocortical dysconnectivity in schizophrenia. Am J Psychiatry. 2012;169(10):1092–1099. doi:10.1176/appi.ajp.2012.12010056

45. Gabernet L, Jadhav SP, Feldman DE, Carandini M, Scanziani M. Somatosensory integration controlled by dynamic thalamocortical feed-forward inhibition. Neuron. 2005;48(2):315–327. doi:10.1016/j.neuron.2005.09.022

46. Courcelles EJ, Kjelsberg K, Convertino L, Nair RR, Witter MP, Nigro MJ. Association cortical areas in the mouse contain a large population of fast-spiking GABAergic neurons that do not express parvalbumin. Eur J Neurosci. 2024;59(12):3236–3255. doi:10.1111/ejn.16341

47. Kim Y, Yang GR, Pradhan K, et al. Brain-wide maps reveal stereotyped cell-type-based cortical architecture and subcortical sexual dimorphism. Cell. 2017;171(2):456–469.e22. doi:10.1016/j.cell.2017.09.020

48. Coutts F, Koutsouleris N, McGuire P. Psychotic disorders as a framework for precision psychiatry. Nat Rev Neurol. 2023;19(4):221–234. doi:10.1038/s41582-023-00779-1

49. Allen P, Luigjes J, Howes OD, et al. Transition to psychosis associated with prefrontal and subcortical dysfunction in ultra high-risk individuals. Schizophr Bull. 2012;38(6):1268–1276. doi:10.1093/schbul/sbr194

50. Modinos G, Allen P, Zugman A, et al. Neural circuitry of novelty salience processing in psychosis risk: association with clinical outcome. Schizophr Bull. 2020;46(3):670–679. doi:10.1093/schbul/sbz089

51. Howes OD, Bose SK, Turkheimer F, et al. Dopamine synthesis capacity before onset of psychosis: a prospective [18F]-DOPA PET imaging study. Am J Psychiatry. 2011;168(12):1311–1317. doi:10.1176/appi.ajp.2011.11010160

52. McBurney-Lin J, Lu J, Zuo Y, Yang H. Locus coeruleus-norepinephrine modulation of sensory processing and perception: A focused review. Neurosci Biobehav Rev. 2019;105:190–199. doi:10.1016/j.neubiorev.2019.06.009

53. Mäki-Marttunen V, Andreassen OA, Espeseth T. The role of norepinephrine in the pathophysiology of schizophrenia. Neurosci Biobehav Rev. 2020;118:298–314. doi:10.1016/j.neubiorev.2020.07.038

54. Nakao K, Singh M, Sapkota K, Fitzgerald A, Hablitz JJ, Nakazawa K. 5-HT2A receptor dysregulation in a schizophrenia relevant mouse model of NMDA receptor hypofunction. Transl Psychiatry. 2022;12(1):168. doi:10.1038/s41398-022-01930-0

55. Ashok AH, Myers J, Reis Marques T, Rabiner EA, Howes OD. Reduced mu opioid receptor availability in schizophrenia revealed with [11C]-carfentanil positron emission tomographic imaging. Nat Commun. 2019;10(1):4493. doi:10.1038/s41467-019-12366-4

56. Lupica CR. Delta and mu enkephalins inhibit spontaneous GABA-mediated IPSCs via a cyclic AMP-independent mechanism in the rat hippocampus. J Neurosci. 1995;15(1 Pt 2):737–749. doi:10.1523/JNEUROSCI.15-01-00737.1995

57. Fusar-Poli P, Borgwardt S, Crescini A, et al. Neuroanatomy of vulnerability to psychosis: A voxel-based meta-analysis. Neuroscience & Biobehavioral Reviews. 2011;35(5):1175–1185. doi:10.1016/j.neubiorev.2010.12.005

58. Lee M, Cernvall M, Borg J, et al. Cognitive function and variability in antipsychotic drug–naive patients with first-episode psychosis: a systematic review and meta-analysis. JAMA Psychiatry. 2024;81(5):468–476. doi:10.1001/jamapsychiatry.2024.0016

