## Supplementary Materials for "Excitation/Inhibition Balance in People at Clinical High Risk for Psychosis: A Functional MRI Study"

### Supplementary Content

**eMethods**

**eTable 1.** Top 20 cortical parcels showing the largest case–control differences in Hurst exponent across PSYSCAN and AMP SCZ samples.

**eTable 2.** Top 20 subcortical parcels showing the largest case–control differences in Hurst exponent across PSYSCAN and AMP SCZ samples.

**eTable 3.** Top 20 cortical and subcortical parcels showing the largest transition-related differences in Hurst exponent in the PSYSCAN.

**eTable 4.** Cox proportional hazards regression coefficients of the retained predictors (parcel-wise Hurst).

**eFigure 1.** Association between head motion and Hurst exponent.

**eFigure 2.** Correspondence between HCPex parcels and the seven functional networks.

**eFigure 3.** Cumulative incidence (Kaplan–Meier failure function) for risk of development of psychosis among CHR-P individuals.

**eFigure 4.** Whole-brain Hurst exponent by antipsychotic medication status in PSYSCAN.

**eFigure 5.** Phenotype effects on whole-brain and network-level Hurst exponent after excluding outliers.

**eFigure 6.** Cross-cohort spatial correspondence of phenotype effects on Hurst exponent.

**eFigure 7.** Phenotype effects on longitudinal trajectories of network-level Hurst exponent.

**eFigure 8.** Transition-related effects on Hurst exponent in the PSYSCAN sample.

**eFigure 9.** Transition-related effects on longitudinal trajectories of network-level Hurst exponent.

**eFigure 10.** Association between phenotype effects on Hurst exponent and cortical thickness across cortical parcels.

**eFigure 11.** Selection of the number of partial least squares components using inner cross-validation.

**eReferences**

**eMethods 1**

**Participant recruitment**

**PSYSCAN**[1]

*General inclusion criteria:*

a) 16-40 years old;

b) Ability to give informed consent (parental/guardian consent is obtained for participants aged <18 years).

*Clinical high risk for psychosis (CHR-P) inclusion criteria:*

a) Inclusion into one of three groups as assessed by the Comprehensive Assessment of At-Risk Mental States (CAARMS version 2006)[2]: i) vulnerability group, ii) attenuated psychosis group, iii) brief intermittent psychosis symptoms group. This instrument has been modified to additionally allow the scoring of the Structured Interview for Psychosis-Risk Syndromes (SIPS).[3] The scoring of the SIPS v.5 is included for comparative purposes and does not constitute inclusion criteria;

b) Inclusion based on meeting criteria for "basic symptoms" which are assessed using the Schizophrenia Proneness Instrument, (SPI-A).[4]

*Healthy control inclusion criteria:*

a) Not having a lifetime history of any DSM-IV Axis-I or Axis-II (borderline, paranoid and schizotypal) disorder;

b) Do not meet CAARMS or SPI-A criteria;

c) No previous use of antipsychotic medication or current use of any psychoactive medication;

d) Do not have a first-degree relative with a lifetime history of affective or nonaffective psychosis (defined by treatment or diagnosis).

*Exclusion criteria:*

a) Any previous neurosurgery or neurological disorder, including epilepsy;

b) History of head injury resulting in unconsciousness lasting at least 1 hour;

c) Pregnancy;

d) Subject is unable to fully comprehend the purpose of the study or make a rational decision whether or not to participate;

e) Estimate of IQ < 70;

f) Antipsychotic medication for > 30 days (cumulative number of days) in the 3 months before the baseline assessments (including self-ratings and screening assessments), at doses that would be adequate for treating a first episode of psychosis (i.e. excludes very low doses);

g) Any past episode of frank psychosis lasting > 7 days;

h) Any contraindications for MRI;

i) Refusing to have blood drawn and/or MRI performed.

**AMP SCZ**[5]

*General inclusion criteria:*

a) Aged 12–30 years;

b) Ability to give informed consent (parental/guardian consent is obtained for participants aged <18 years);

c) Meeting either CHR-P or healthy control criteria.

*Clinical high risk for psychosis (CHR-P) inclusion criteria:*

a) Meet CAARMS-defined (Trait Vulnerability; Attenuated Psychotic Symptoms; Brief Limited Intermittent Psychotic Symptoms) or Structured Interview for Prodromal Syndromes-defined (Brief Intermittent Psychotic Syndrome Current Progression; Attenuated Positive Symptom Syndrome Current Progression; Genetic Risk and Deterioration Current Progression) diagnostic criteria for CHR determined using a newly developed instrument, the Positive SYmptoms and Diagnostic Criteria for the CAARMS Harmonized with the SIPS (PSYCHS).[6]

*Healthy control inclusion criteria:*

a) Do not meet CHR-P criteria or have a current or past Cluster A personality disorder;

b) Not receiving any current treatment with psychotropic medication;

c) Do not have a family history (in first-degree relatives) of psychotic spectrum disorders.

*Exclusion criteria:*

a) Antipsychotic medication exposure equivalent to a total lifetime haloperidol dose of >50 mg, estimated based on available information, or current antipsychotic medication at time of baseline assessment;

b) Documented history of intellectual disability;

c) Past or current clinically relevant central nervous system disorder;

d) Traumatic brain injury rated 7 or above on the Traumatic Brain Injury screening instrument;

e) Current or past psychotic disorder.

**eMethods 2**

**<u>Imaging parameters</u>**

**PSYSCAN**[1]

MRI data were acquired on 3T Siemens TrioTim, Siemens Verio, Philips Achieva, and GE Signa scanners. Participants were instructed to remain still for 9 minutes. To minimize inter-site variability, all PSYSCAN consortium sites implemented a harmonized imaging acquisition protocol and completed a site qualification procedure to ensure that the standardized protocol could be applied locally. $T_1$-weighted ($T_1$-w) images parameters: voxel size = 1.2 × 1.0 × 1.0 mm$^3$, 176 slices (thickness = 1.2 mm), repetition time (TR) = 2300 ms, echo time (TE) = 2.96 ms, flip angle = 9°, inversion time = 900 ms, field of view = 256 × 240 mm$^2$, and acquisition matrix = 256 × 240. Resting-state functional MRI (rsfMRI) parameters: voxel size = 3.3 mm, 240 volumes, 48 slices (thickness = 3.0 mm), TR = 2500 ms, TE = 30 ms, flip angle = 80°, field of view = 211 × 211 $mm^2$ and matrix = 64 × 64.

**AMP SCZ**[5]

MRI data were acquired on 3T Siemens Prisma, Siemens Skyra, Siemens Vida, General Electric MR750, and Philips Achieva DDAS scanners. Participants completed four 5-minute runs, consisting of two anterior-to-posterior and two posterior-to-anterior phase-encoding runs. The structural and rsfMRI scans are well harmonized regarding scan parameters across vendors and platforms. $T_1$-w images parameters: voxel size = 0.8 mm, 208 slices (thickness = 0.8 mm), TR = 2500 ms, TE = 2.07 ms, flip angle = 8°, inversion time = 1000 ms, field of view = 256 × 240 $mm^2$, and acquisition matrix = 320 × 300. rsfMRI scans parameters: voxel size = 2.4 mm, 333–343 volumes per run, 60 slices (thickness = 2.4 mm), TR = 900 ms, TE = 35 ms, flip angle = 52°, multiband factor = 6, field of view = 216 × 216 $mm^2$ and matrix = 90 × 90. Further details are provided in the AMP SCZ imaging protocol paper.[7]

**Imaging preprocessing**

Neuroimaging data preprocessing was conducted using fMRIPrep (version 23.2.0).[8] For T1-w structural MRI, preprocessing included: (i) correction for intensity non-uniformity using N4BiasFieldCorrection[9]; (ii) skull stripping using the ANTs-based antsBrainExtraction.sh workflow[10]; (iii) tissue segmentation into gray matter (GM), white matter (WM), and cerebrospinal fluid (CSF) using FAST (FSL)[11]; (iv) nonlinear spatial normalization to the MNI152NLin2009cAsym standard space using ANTs antsRegistration[10]; (v) Surface reconstruction was performed using the FreeSurfer via the recon-all command.[12]

Core preprocessing of rsfMRI data included: (i) generation of a reference volume and its skull-stripped version; (ii) slice-timing correction using AFNI 3dTShift[13]; (iii) rigid-body head-motion correction using FSL MCFLIRT,[11] with computation of framewise displacement (FD) and DVARS[14]; (iv) co-registration of the BOLD reference image to the skull-stripped T1-w structural image using bbregister (FreeSurfer)[15]; and (v) spatial normalization to the MNI152NLin2009cAsym standard space. Denoising steps included: (i) spatial smoothing with a 6 mm FWHM Gaussian kernel using AFNI 3dBlurInMask[13]; (ii) intensity normalization to a whole-brain median of 1000 using AFNI 3dBrickStat and 3dcalc[13]; (iii) wavelet-based time-series despiking using the BrainWavelet toolbox[16]; (iv) 13-parameter confound regression including six motion parameters, their first-order temporal derivatives, and ventricular cerebrospinal fluid (CSF) signal using AFNI 3dBandpass with the -ort option; no temporal bandpass filtering was applied.[13]

Wavelet-based denoising was used because it can reduce motion-related artifacts that are spatially and temporally heterogeneous, including both linear and non-linear effects, without requiring data scrubbing.[16] This was particularly important for the present study, as our time-series based analyses relied on preserving the temporal continuity of the signal. Data scrubbing was not appropriate because removing volumes disrupts the temporal structure of the fMRI time series and could therefore bias estimation of the Hurst exponent (HE), which is intended to capture long-memory properties of the signal.

**Quality checks**

All preprocessing outputs underwent visual quality inspection to ensure accurate brain extraction, tissue segmentation, and functional-to-anatomical alignment. For denoising, we evaluated wavelet despiking by extracting the whole-brain mean time series before and after despiking, visually inspecting the reduction of transient spikes, and summarizing spike percentages across voxels. The effectiveness of nuisance regression was assessed by calculating the mean voxelwise correlation between the denoised fMRI data and the confound regressors, with values expected to approach zero. Finally, as a biologically meaningful validation step, posterior cingulate cortex seed-based functional connectivity analysis was performed to verify that the resulting connectivity pattern was consistent with the expected default mode network topology.[17]

**eMethods 3**

**Hurst exponent estimation**

Consistent with prior HE work,[18] participants with a mean FD >0.4 mm were excluded. For AMP SCZ, mean FD was averaged across available runs within each participant/session before applying this threshold. The mean FD criterion excluded 28 participants from PSYSCAN and 15 from AMP SCZ. Sites with fewer than 10 participants were also excluded, excluding 4 sites with 15 participants from PSYSCAN and 7 sites with 38 participants from AMP SCZ.

Regional HE was estimated using the bfn_mfin_ml algorithm from the nonfractal toolbox (https://github.com/wonsang/nonfractal), which implements a fractionally integrated process (FIP) model with maximum likelihood estimation. Unlike the fractional Gaussian noise (fGn) model, which assumes stationarity and restricts HE to values below 1, the FIP model removes the stationarity assumption and allows HE estimates to exceed 1. This greater flexibility avoids the ceiling effects associated with the fGn model and may improve the characterization of interindividual differences in temporal dynamics, particularly when HE values approach or exceed 1. Within the FIP framework, HE values above 1 indicate non-stationary time series with long-memory characteristics.[18] The "filter" parameter was set to "haar," and the "ub" (upper bound) and "lb" (lower bound) were specified as [1.5, 10] and [-0.5, 0], consistent with prior HE studies.[18,19] For AMP SCZ, parcel-wise HE estimates were averaged across all runs to obtain a single HE estimate per parcel for each participant/session.

To assess whether head motion was associated with HE, we examined the relationship between whole-brain mean HE and mean FD. An ordinary least squares regression (OLS) model was fitted across participants, with whole-brain mean HE as the dependent variable, mean FD as the predictor, and phenotype included as a covariate (HE ~ mean FD + phenotype) (**eFigure 1**).

### Site harmonization

In PSYSCAN,[1] six healthy subjects were scanned across six sites that contributed the majority of scans (Melbourne, Amsterdam, Maastricht, London, Toronto, and Seoul; 84.6% of all scans) as travelling heads. Site-specific offsets were then estimated using travelling heads data via linear mixed-effects models with random intercepts for subject and site. The offsets were subtracted for all subjects at corresponding sites. The two remaining sites (Naples and Madrid) without travelling-subject data were further harmonized using neuroComBat,[20] with site as the batch variable and age, sex, phenotype/transition status, MeanFD as covariates. Travelling heads sites were considered a single mega-site following mean-offset correction. In AMP SCZ, travelling-subject data were not available, so data from 31 sites were harmonized using the neuroComBat[20] with same covariates. All downstream group-level analyses were performed on the harmonized data.

### Network assignment

To map the HCPex atlas[21] onto the functional networks, the Yeo 7-network atlas[22] in MNI152 1 mm space was resampled to the HCPex atlas space using nearest-neighbor interpolation to preserve the discrete network labels (**eFigure 2**). Voxel-wise overlap between each HCPex parcel and the resampled Yeo atlas was then quantified. For each parcel, the proportion of voxels assigned to each of the seven Yeo networks was calculated, generating a parcel-by-network proportion matrix for weighted network-level analyses.

Network-level HE values were subsequently computed using the parcel-by-network proportion matrix as soft assignment weights. For each subject and each Yeo 7 network, a weighted mean HE value was defined as:

$$\text{NetworkValue}_{s,n} = \frac{\sum_i w_{i,n} H_{s,i}}{\sum_i w_{i,n}}$$

where $H_{s,i}$ represents the HE value of parcel $i$ in subject $s$, and $w_{i,n}$ represents the proportional overlap between parcel $i$ and network $n$. Accordingly, parcels were allowed to contribute to multiple networks in proportion to their overlap, rather than being assigned to a single network only.

**eMethods 4**

**Cox proportional hazards regression for psychosis transition prediction**

Follow-up time was censored at 3 years. The cumulative incidence of transition to psychosis was visualized using the Kaplan–Meier failure function (1 − survival)[23] with Greenwood 95% confidence intervals (CIs)[24] (**eFigure 3**). Cox proportional hazards regression with elastic net regularisation was used to develop the transition prediction model for participants with varying follow-up durations and high-dimensional predictors.[25] The caret R package (version 7.0.1) was used to generate cross-validation (CV) folds stratified by transition status, while penalised Cox models were fitted using the glmnet R package, with survival outcomes specified using follow-up time and transition status. The analysis comprised two stages: (i) a primary repeated nested CV procedure, with three inner folds repeated five times for hyperparameter tuning and three outer folds repeated five times for performance estimation on unseen data; and (ii) a final post hoc model fitted to the complete dataset for coefficient estimation and interpretation.

For the primary CV framework, each outer repetition randomly divided the data into 3 outer folds, with 2 folds used as the outer training set and the remaining fold used for model testing. Missing HE values and covariates required for harmonization were imputed within the CV framework using random forest imputation (missForest R package, version 1.5; default parameters), with multivariate imputation by chained equations (mice R package, version 3.18.0; default parameters) used as a fallback. With each outer split, neuroComBat harmonization was fitted using the outer training data only, using the same batch variable and covariates described above, and the estimated harmonization parameters were subsequently applied to the corresponding held-out test fold. Parcel-wise HE values were residualised for age, sex, and MeanFD and subsequently standardised using parameters estimated from the outer training set, with the same transformations applied to the corresponding test fold. Predictors with near-zero variance were removed.

The preprocessed outer training set was further divided into 3 inner folds, repeated 5 times, to select the elastic net hyperparameters: the mixing parameters (α; balancing LASSO and ridge penalties) and regularisation parameters (λ; determining the degree of coefficient shrinkage). The tuning grid comprised α values from 0.1 to 0.9 and 100 logarithmically spaced λ values ranging from 1 to 0.001. For each α–λ combination, repeated inner CV generated out-of-fold (OOF) predictions, and the combination with the highest mean C-index was selected. The selected model was then refitted to the complete outer training set, and its performance was evaluated on the held-out outer test folds. The three-fold outer CV procedure was repeated five times using different fold allocations to improve the robustness.

Model performance was evaluated across three domains. First, discrimination was assessed using Harrell's C-index, which reflects the model's ability to distinguish individuals at higher versus lower risk of transition, with values above 0.5 indicating better-than-chance discrimination. Second, threshold-dependent classification performance, including balanced accuracy (BAC), sensitivity and specificity, was assessed across cutoffs from 0.00 to 1.00 in increments of 0.01. The cutoff yielding the highest BAC was used to calculate the final classification metrics. Third, overall predictive accuracy was assessed using the Brier score, with values closer to 0 indicating better performance and values above 0.25 generally indicating a poor model. Metrics were presented as mean and 95% confidence interval across all outer CV folds and repeats.

For interpretability, a final post hoc elastic-net Cox model was tuned using three-fold CV repeated five times on the full dataset and then refitted to obtain coefficients. These coefficients were reported to characterise the regional HE features contributing to prediction, whereas repeated nested CV estimates were used as the primary measures of model performance.

**eMethods 5**

**Cortical thickness (CT) calculation and analysis**

CT was estimated by FreeSurfer at each vertex in native surface space based on the reconstructed white and pial surfaces.[26] The resulting native-space thickness maps were subsequently resampled to the fsaverage surface. For regional analyses, the fsaverage thickness maps from each hemisphere were parcellated using the HCPex atlas,[21] and mean CT was calculated for each cortical parcel. This procedure yielded a subject-by-parcel CT matrix for subsequent analyses.

Scanner-related variability was addressed using the same harmonization strategy as for HE: a combined travelling-head plus neuroCombat[20] approach in PSYSCAN, and neuroCombat alone in AMP-SCZ, where no travelling-subject data were available. All subsequent analyses were conducted on the harmonized CT measures.

For each cortical parcel, an ordinary least squares (OLS) model was fitted according to the formula $Y = \beta_0 + \beta_1\text{Phenotype} + \beta_2\text{Age} + \beta_3\text{Sex} + \beta_4\text{MeanFD} + \varepsilon$. The regression coefficient for the Phenotype term ($\beta_1$) was extracted to generate parcel-wise maps of phenotype-associated effects on CT. Spatial correspondence between the CT $\beta$-map and the HE $\beta$-map was then quantified using Pearson correlation. Statistical significance was assessed using a non-parametric permutation test with 10,000 iterations, in which phenotype labels were randomly shuffled and the full set of models was refitted at each iteration.

**Partial Least Square Regression (PLSR) pipeline for receptor density map analysis**

PLSR was used as a multivariate association method to relate high-dimensional parcel-wise receptor density data to phenotype-associated variation in HE. The parcel-wise outcome variable was defined as the phenotype regression coefficient ($\beta$) estimated separately for each parcel from an OLS model of the form: $HE = \beta_0 + \beta_1\text{Phenotype} + \beta_2\text{Age} + \beta_3\text{Sex} + \beta_4\text{MeanFD} + \varepsilon$. Nineteen neurotransmitter receptor density maps compiled by Hansen et al[27] were used as predictors (https://github.com/netneurolab/hansen_receptors). The receptor maps included serotonin (5-$HT_{1A}$, 5-$HT_{1B}$, 5-$HT_{2A}$, 5-$HT_4$, 5-$HT_6$, and 5-HTT), dopamine ($D_1$, $D_2$, and DAT), norepinephrine (NET), histamine ($H_3$), acetylcholine ($\alpha_4\beta_2$, $M_1$, and VAChT), cannabinoid ($CB_1$), opioid (MOR), glutamate (NMDA and $mGluR_5$), and GABA ($GABA_{A/BZ}$). Volumetric PET/SPECT maps in MNI152 space were parcellated to the HCPex[21] cortical atlas using neuromaps' Parcellater.[28] Where multiple tracers were available for a given target, maps were z-scored and combined using sample-size-weighted averaging to yield a single parcel-wise map per receptor. The resulting predictor matrix ($X$) comprised parcel-wise receptor densities, and the response vector ($Y$) comprised the corresponding parcel-wise phenotype-associated $\beta$ estimates.

PLSR was evaluated under cross-validation (CV) to guide model complexity selection and quantify out-of-sample association strength. The analysis pipeline comprised two stages: (i) a primary nested CV model used to assess the statistical significance and out-of-sample generalisability of the receptor–HE association while selecting the number of latent variables (LVs), and (ii) a final parsimonious model with a fixed number of LVs, informed by the primary analysis, used for interpretation and Variable Importance in Projection (VIP) estimation. In both stages, model performance was quantified using cross-validated $R^2$ and Pearson's $r$ between observed and predicted parcel-wise $\beta$ values. Predictors and outcomes were scaled within each CV fold using parameters estimated from the training data only, and these parameters were then applied to the corresponding held-out fold.

For the primary analysis, nested CV was implemented using an outer 10-fold loop and an inner 5-fold loop.[29] Within each outer training set, candidate models with different numbers of LVs were evaluated using inner-loop $R^2$. A one-standard-error (1-SE) rule[30] was applied within each outer training set to select the smallest LV dimensionality whose mean inner-CV performance fell within one standard error of the best-performing model (**eFigure 11**). This procedure generated out-of-fold (OOF) predictions across the 10 outer test folds; overall cross-validated $R^2$ and Pearson's $r$ were then calculated by comparing the pooled OOF predictions with the observed parcel-wise β values. In parallel, inner-CV performance curves were aggregated across outer folds to derive a global 1-SE solution, which was used to define the final parsimonious model. Statistical significance of the primary model was assessed using nonparametric permutation testing ($n = 10{,}000$). For each permutation, phenotype labels were randomly shuffled across subjects, parcel-wise OLS models were re-estimated to generate a permuted phenotype-effect map, and the full primary PLSR pipeline was repeated using this permuted $Y$ vector. This produced null distributions for cross-validated $R^2$ and Pearson's $r$, from which empirical permutation $p$-values were computed.

The parsimonious model was then refit using the selected LV and the same 10-fold CV partitioning, but without further nested hyperparameter optimisation. This fixed-complexity refit was used to characterise the dominant and stable covariance structure underlying the receptor–HE association. VIP scores were computed from this parsimonious model to quantify the contribution of individual receptor predictors to the latent structure identified in the primary analysis. VIP stability was further assessed by bootstrap resampling of parcels (B=10,000). Receptors with mean VIP values > 1 were considered to contribute above average to the model.

**eTable 1.** Top 20 cortical parcels showing the largest case–control differences in Hurst exponent across PSYSCAN and AMP SCZ samples.

| Parcel | β | t | p | q |
|---|---|---|---|---|
| ***PSYSCAN sample*** | | | | |
| EarlyAuditory_LBelt_Left | -0.058 | -3.528 | <0.001 | 0.082 |
| VentralStreamVisual_V8_Left | -0.058 | -4.003 | <0.001 | 0.028 |
| InsularandFrontalOpercular_MI_Left | -0.052 | -3.261 | 0.001 | 0.082 |
| EarlyAuditory_A1_Left | -0.051 | -3.259 | 0.001 | 0.082 |
| InsularandFrontalOpercular_PoI2_Left | -0.051 | -3.006 | 0.003 | 0.086 |
| PosteriorOpercular_OP2-3_Left | -0.049 | -3.161 | 0.002 | 0.082 |
| VentralStreamVisual_V8_Right | -0.049 | -3.155 | 0.002 | 0.082 |
| PosteriorOpercular_OP1_Right | -0.048 | -2.891 | 0.004 | 0.098 |
| ParacentralLobularandMidCingulate_6ma_Left | -0.047 | -3.059 | 0.002 | 0.086 |
| PrimaryVisual_V1_Left | -0.047 | -3.017 | 0.003 | 0.086 |
| EarlyAuditory_PBelt_Left | -0.046 | -3.144 | 0.002 | 0.082 |
| EarlyVisual_V3_Right | -0.046 | -2.967 | 0.003 | 0.090 |
| InsularandFrontalOpercular_PoI1_Left | -0.046 | -2.852 | 0.005 | 0.098 |
| SomatosensoryandMotor_1_Right | -0.045 | -2.786 | 0.006 | 0.113 |
| PosteriorCingulate_ProS_Left | -0.045 | -3.158 | 0.002 | 0.082 |
| EarlyVisual_V4_Right | -0.045 | -2.854 | 0.005 | 0.098 |
| PosteriorOpercular_OP4_Left | -0.042 | -3.104 | 0.002 | 0.083 |
| LateralTemporal_TE1p_Left | -0.040 | -2.764 | 0.006 | 0.114 |
| PosteriorCingulate_RSC_Left | -0.038 | -2.909 | 0.004 | 0.098 |
| AuditoryAssociation_A4_Left | -0.036 | -2.709 | 0.007 | 0.121 |
| ***AMP SCZ sample*** | | | | |
| PosteriorOpercular_OP1_Right | -0.029 | -3.909 | <0.001 | 0.014 |
| PosteriorOpercular_OP2-3_Right | -0.029 | -3.866 | <0.001 | 0.014 |
| PosteriorOpercular_OP4_Right | -0.024 | -3.896 | <0.001 | 0.014 |
| EarlyAuditory_MBelt_Right | -0.023 | -3.250 | 0.001 | 0.034 |
| SomatosensoryandMotor_1_Right | -0.023 | -3.552 | <0.001 | 0.029 |
| EarlyAuditory_A1_Right | -0.022 | -3.199 | 0.001 | 0.036 |
| PosteriorCingulate_ProS_Right | -0.022 | -3.240 | 0.001 | 0.034 |
| PosteriorOpercular_OP4_Left | -0.022 | -3.482 | 0.001 | 0.031 |
| ParacentralLobularandMidCingulate_5mv_Left | -0.022 | -3.640 | <0.001 | 0.026 |
| ParacentralLobularandMidCingulate_5m_Left | -0.021 | -3.032 | 0.002 | 0.045 |
| SuperiorParietal_7AL_Right | -0.021 | -3.440 | 0.001 | 0.031 |
| EarlyAuditory_PFcm_Left | -0.021 | -3.371 | 0.001 | 0.034 |
| AuditoryAssociation_STGa_Right | -0.020 | -3.095 | 0.002 | 0.043 |
| ParacentralLobularandMidCingulate_5L_Left | -0.020 | -3.185 | 0.001 | 0.036 |
| EarlyAuditory_PFcm_Right | -0.020 | -3.241 | 0.001 | 0.034 |
| AnteriorCingulateandMedialPrefrontal_pOFC_Right | -0.020 | -3.290 | 0.001 | 0.034 |
| PosteriorOpercular_43_Right | -0.019 | -3.318 | 0.001 | 0.034 |
| AuditoryAssociation_TA2_Left | -0.019 | -3.076 | 0.002 | 0.043 |
| PosteriorOpercular_FOP1_Left | -0.019 | -3.031 | 0.002 | 0.045 |
| ParacentralLobularandMidCingulate_23c_Right | -0.017 | -3.150 | 0.002 | 0.038 |

***Notes*:** Results are based on parcel-wise ordinary least squares (OLS) regression models: $HE = \beta_0 + \beta_1 Phenotype + \beta_2 Age + \beta_3 Sex + \beta_4 MeanFD + \varepsilon$, where the Phenotype term represents the clinical high risk for psychosis (CHR-P) vs. healthy controls (HC) contrast. Columns report the regression coefficient (β), corresponding t-value, nominal p-value, and FDR-corrected q-value. Negative β values indicate lower Hurst exponent in CHR-P individuals relative to HCs.

**eTable 2.** Top 20 subcortical parcels showing the largest case–control differences in Hurst exponent across PSYSCAN and AMP SCZ samples.

| Parcel | β | t | p | q |
|---|---|---|---|---|
| ***PSYSCAN sample*** | | | | |
| Thalamic_Nuclei_MDl_Left | -0.047 | -2.558 | 0.011 | 0.432 |
| Thalamic_Nuclei_MDm_Right | -0.043 | -2.393 | 0.017 | 0.432 |
| Thalamic_Nuclei_PuM_Right | -0.042 | -2.328 | 0.021 | 0.432 |
| Thalamic_Nuclei_PuA_Left | -0.033 | -1.955 | 0.051 | 0.540 |
| Thalamic_Nuclei_VLp_Left | -0.033 | -2.009 | 0.045 | 0.540 |
| Thalamic_Nuclei_VPL_Left | -0.032 | -1.959 | 0.051 | 0.540 |
| Subcortical_Region_Putam_Right | -0.029 | -1.846 | 0.066 | 0.555 |
| Thalamic_Nuclei_CeM_Right | -0.028 | -1.712 | 0.088 | 0.555 |
| Subcortical_Region_Caud_Left | -0.028 | -1.751 | 0.081 | 0.555 |
| Subcortical_Region_Gpe_Right | -0.027 | -1.737 | 0.083 | 0.555 |
| Thalamic_Nuclei_MDl_Right | -0.025 | -1.559 | 0.120 | 0.620 |
| Thalamic_Nuclei_PuM_Left | -0.025 | -1.388 | 0.166 | 0.620 |
| Thalamic_Nuclei_VLa_Left | -0.025 | -1.578 | 0.116 | 0.620 |
| Thalamic_Nuclei_MDm_Left | -0.025 | -1.384 | 0.167 | 0.620 |
| Thalamic_Nuclei_CM_Left | -0.023 | -1.417 | 0.158 | 0.620 |
| Subcortical_Region_Gpi_Right | -0.023 | -1.500 | 0.135 | 0.620 |
| Subcortical_Region_Caud_Right | -0.022 | -1.404 | 0.161 | 0.620 |
| Thalamic_Nuclei_PuA_Right | -0.020 | -1.229 | 0.220 | 0.709 |
| Thalamic_Nuclei_VA_Left | -0.019 | -1.132 | 0.259 | 0.709 |
| Thalamic_Nuclei_VPL_Right | -0.019 | -1.166 | 0.245 | 0.709 |
| ***AMP SCZ sample*** | | | | |
| Thalamic_Nuclei_MV(Re)_Left | -0.023 | -3.852 | <0.001 | 0.008 |
| Thalamic_Nuclei_MV(Re)_Right | -0.020 | -3.262 | 0.001 | 0.010 |
| Subcortical_Region_Caud_Left | -0.019 | -3.518 | <0.001 | 0.010 |
| Thalamic_Nuclei_VLa_Right | -0.019 | -2.912 | 0.004 | 0.026 |
| Subcortical_Region_Caud_Right | -0.018 | -3.332 | <0.001 | 0.010 |
| Thalamic_Nuclei_AV_Left | -0.017 | -3.425 | <0.001 | 0.010 |
| Thalamic_Nuclei_VLa_Left | -0.016 | -2.419 | 0.016 | 0.071 |
| Subcortical_Region_VTA_Right | -0.016 | -3.312 | <0.001 | 0.010 |
| Subcortical_Region_VTA_Left | -0.016 | -3.310 | <0.001 | 0.010 |
| Thalamic_Nuclei_Pf_Left | -0.015 | -2.988 | 0.003 | 0.023 |
| Thalamic_Nuclei_CeM_Left | -0.014 | -2.568 | 0.010 | 0.065 |
| Thalamic_Nuclei_LGN_Right | -0.014 | -2.297 | 0.022 | 0.086 |
| Subcortical_Region_NAc_Right | -0.013 | -2.519 | 0.012 | 0.065 |
| Thalamic_Nuclei_VLp_Right | -0.012 | -1.927 | 0.054 | 0.137 |
| Subcortical_Region_SNpc_Right | -0.012 | -2.500 | 0.013 | 0.065 |
| Thalamic_Nuclei_VLp_Left | -0.012 | -1.965 | 0.050 | 0.136 |
| Thalamic_Nuclei_PuL_Right | -0.012 | -1.939 | 0.053 | 0.137 |
| Subcortical_Region_NAc_Left | -0.012 | -2.203 | 0.028 | 0.097 |
| Subcortical_Region_Septum_Left | -0.011 | -2.479 | 0.013 | 0.065 |
| Subcortical_Region_SNpc_Left | -0.011 | -2.392 | 0.017 | 0.071 |

***Notes***: Results are based on parcel-wise ordinary least squares (OLS) regression models: HE = $\beta_0 + \beta_1$Phenotype + $\beta_2$Age + $\beta_3$Sex + $\beta_4$MeanFD + ε, where the Phenotype term represents the clinical high risk for psychosis (CHR-P) vs. healthy controls (HC) contrast. Columns report the regression coefficient (β), corresponding t-value, nominal p-value, and FDR-corrected q-value. Negative β values indicate lower Hurst exponent in CHR-P individuals relative to HCs.

**eTable 3.** Top 20 cortical and subcortical parcels showing the largest transition-related differences in Hurst exponent in the PSYSCAN sample.

| Parcel | β | t | p | q |
|---|---|---|---|---|
| ***Cortical parcels*** | | | | |
| AnteriorCingulateandMedialPrefrontal_a24_Right | 0.090 | 3.019 | 0.003 | 0.998 |
| AnteriorCingulateandMedialPrefrontal_25_Right | 0.073 | 2.282 | 0.024 | 0.998 |
| AnteriorCingulateandMedialPrefrontal_a24_Left | 0.073 | 2.512 | 0.013 | 0.998 |
| EarlyAuditory_MBelt_Left | 0.071 | 2.042 | 0.043 | 0.998 |
| AnteriorCingulateandMedialPrefrontal_25_Left | 0.071 | 2.262 | 0.025 | 0.998 |
| MedialTemporal_PreS_Right | 0.069 | 2.087 | 0.038 | 0.998 |
| ParacentralLobularandMidCingulate_5mv_Right | 0.063 | 2.166 | 0.032 | 0.998 |
| DorsolateralPrefrontal_9a_Left | 0.061 | 2.081 | 0.039 | 0.998 |
| SuperiorParietal_7PL_Right | 0.059 | 2.102 | 0.037 | 0.998 |
| AuditoryAssociation_STGa_Left | -0.058 | -1.928 | 0.055 | 0.998 |
| AnteriorCingulateandMedialPrefrontal_p32_Right | 0.057 | 2.101 | 0.037 | 0.998 |
| AnteriorCingulateandMedialPrefrontal_p24_Left | 0.056 | 1.801 | 0.073 | 0.998 |
| DorsalStreamVisual_V7_Left | -0.055 | -1.901 | 0.059 | 0.998 |
| PosteriorCingulate_POS1_Right | 0.055 | 1.868 | 0.063 | 0.998 |
| InferiorParietal_PF_Right | 0.053 | 1.905 | 0.058 | 0.998 |
| PosteriorOpercular_43_Left | 0.051 | 1.944 | 0.053 | 0.998 |
| ParacentralLobularandMidCingulate_23c_Right | 0.051 | 1.793 | 0.075 | 0.998 |
| PosteriorOpercular_FOP1_Left | 0.050 | 1.794 | 0.075 | 0.998 |
| AnteriorCingulateandMedialPrefrontal_10r_Left | 0.049 | 1.987 | 0.048 | 0.998 |
| PosteriorCingulate_POS2_Left | 0.047 | 1.936 | 0.054 | 0.998 |
| ***Subcortical parcels*** | | | | |
| Subcortical_Region_SNpc_Right | 0.078 | 2.840 | 0.005 | 0.316 |
| Subcortical_Region_VTA_Right | 0.063 | 2.100 | 0.037 | 0.625 |
| Thalamic_Nuclei_VPL_Right | 0.061 | 1.940 | 0.054 | 0.625 |
| Subcortical_Region_SNpr_Right | 0.060 | 1.896 | 0.060 | 0.625 |
| Thalamic_Nuclei_CeM_Right | 0.060 | 1.923 | 0.056 | 0.625 |
| Subcortical_Region_Putam_Left | 0.055 | 1.688 | 0.093 | 0.692 |
| Subcortical_Region_Gpi_Right | 0.054 | 1.904 | 0.058 | 0.625 |
| Thalamic_Nuclei_PuM_Right | 0.053 | 1.545 | 0.124 | 0.692 |
| Subcortical_Region_Caud_Right | 0.048 | 1.588 | 0.114 | 0.692 |
| Thalamic_Nuclei_PuM_Left | 0.047 | 1.343 | 0.181 | 0.713 |
| Thalamic_Nuclei_L-Sg_Right | 0.047 | 1.606 | 0.110 | 0.692 |
| Thalamic_Nuclei_VA_Right | 0.046 | 1.514 | 0.132 | 0.692 |
| Subcortical_Region_Nb_Right | 0.044 | 1.521 | 0.130 | 0.692 |
| Subcortical_Region_Putam_Right | 0.043 | 1.364 | 0.174 | 0.713 |
| Subcortical_Region_SNpc_Left | 0.043 | 1.373 | 0.172 | 0.713 |
| Thalamic_Nuclei_VLp_Right | 0.041 | 1.356 | 0.177 | 0.713 |
| Subcortical_Region_MB_Left | 0.041 | 1.145 | 0.254 | 0.721 |

| | | | | |
|---|---|---|---|---|
| Thalamic_Nuclei_Pf_Left | 0.038 | 1.139 | 0.256 | 0.721 |
| Subcortical_Region_NAc_Left | 0.036 | 1.194 | 0.234 | 0.721 |
| Thalamic_Nuclei_LGN_Left | 0.036 | 1.152 | 0.251 | 0.721 |

***Notes***: Results are based on parcel-wise ordinary least squares (OLS) regression models: $HE = \beta_0 + \beta_1 Phenotype + \beta_2 Age + \beta_3 Sex + \beta_4 MeanFD + \varepsilon$, where the Phenotype term represents the clinical high risk for psychosis individuals who later transitioned to psychosis (CHR-T) vs. those who did not transition (CHR-NT). Columns report the regression coefficient (β), corresponding t-value, nominal p-value, and FDR-corrected q-value. Negative β values indicate lower Hurst exponent in CHR-T individuals relative to CHR-NT.

**eTable 4.** Cox proportional hazards regression coefficients of the retained predictors (parcel-wise Hurst).

| Parcel | Coefficient_standardized |
|---|---|
| ***Cortical parcels*** | |
| AnteriorCingulateandMedialPrefrontal_10r_Left | 0.156 |
| AnteriorCingulateandMedialPrefrontal_33pr_Right | -0.121 |
| AnteriorCingulateandMedialPrefrontal_a24_Left | 0.069 |
| AnteriorCingulateandMedialPrefrontal_a24_Right | 0.382 |
| AnteriorCingulateandMedialPrefrontal_p24_Right | 0.011 |
| AnteriorCingulateandMedialPrefrontal_pOFC_Left | 0.015 |
| AuditoryAssociation_A4_Left | 0.089 |
| AuditoryAssociation_STGa_Left | -0.221 |
| DorsalStreamVisual_V3B_Left | -0.160 |
| DorsalStreamVisual_V3B_Right | -0.040 |
| DorsolateralPrefrontal_8Av_Right | 0.041 |
| DorsolateralPrefrontal_SFL_Left | -0.042 |
| EarlyAuditory_A1_Left | -0.006 |
| EarlyAuditory_MBelt_Left | 0.034 |
| EarlyAuditory_PBelt_Right | -0.139 |
| InferiorParietal_PF_Left | 0.036 |
| InferiorParietal_PF_Right | 0.228 |
| InsularandFrontalOpercular_FOP5_Right | -0.009 |
| MT+ComplexandNeighboringVisualAreas_V4t_Left | -0.052 |
| MedialTemporal_PHA1_Right | 0.000 |
| MedialTemporal_PHA3_Right | 0.006 |
| MedialTemporal_PeEc_Left | 0.014 |
| OrbitalandPolarFrontal_10pp_Right | -0.284 |
| ParacentralLobularandMidCingulate_5mv_Right | 0.187 |
| PosteriorCingulate_POS2_Left | 0.091 |
| Premotor_PEF_Left | -0.132 |
| SuperiorParietal_7PL_Right | 0.167 |
| VentralStreamVisual_VVC_Right | 0.060 |
| ***Subcortical parcels*** | |
| Subcortical_Region_SNpc_Right | 0.320 |
| Subcortical_Region_SNpr_Right | 0.011 |
| Subcortical_Region_Gpi_Right | 0.066 |

***Notes:*** Coefficients are standardized coefficients from the final post hoc elastic-net Cox proportional hazards model fitted to the full PSYSCAN CHR-P sample. Only predictors with non-zero coefficients are shown.

Positive coefficients indicate that higher regional Hurst exponent values were associated with a higher hazard of transition to psychosis, whereas negative coefficients indicate an association with a lower hazard.

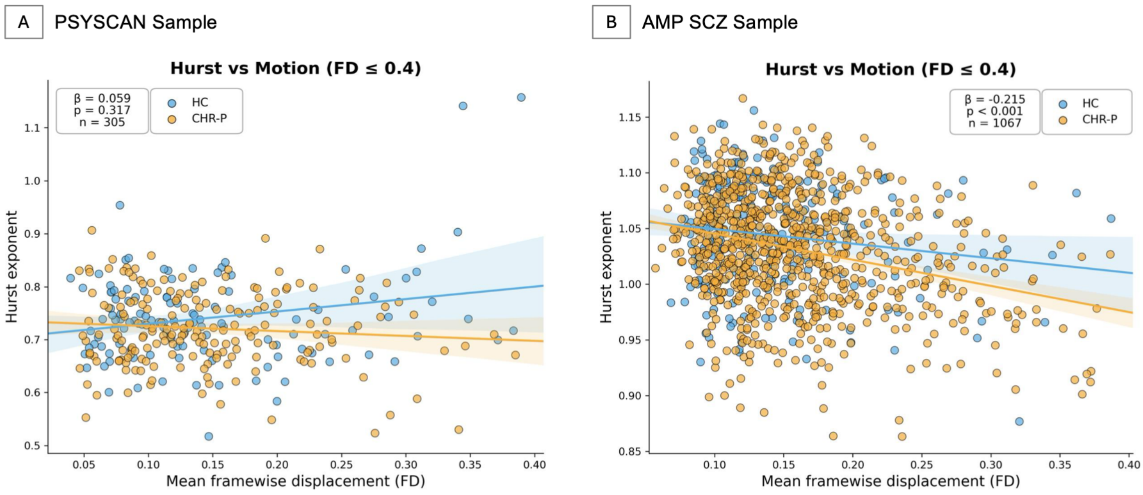


**eFigure 1. Association between head motion and Hurst exponent.** Scatter plots show the association between mean framewise displacement (MeanFD) and whole-brain mean Hurst exponent (HE) in A) the PSYSCAN sample and B) the AMP SCZ sample. Each point represents one participant, with healthy controls (HCs) shown in blue and clinical high risk for psychosis (CHR-P) participants shown in orange. Solid lines represent group-specific linear regression fits for visualization, with shaded areas indicating 95% confidence intervals. In PSYSCAN, mean FD was not significantly associated with HE ($\beta = 0.059$, $p = 0.317$, $n = 305$), whereas in AMP SCZ, higher mean FD was associated with lower HE ($\beta = -0.215$, $p < 0.001$, $n = 1067$).

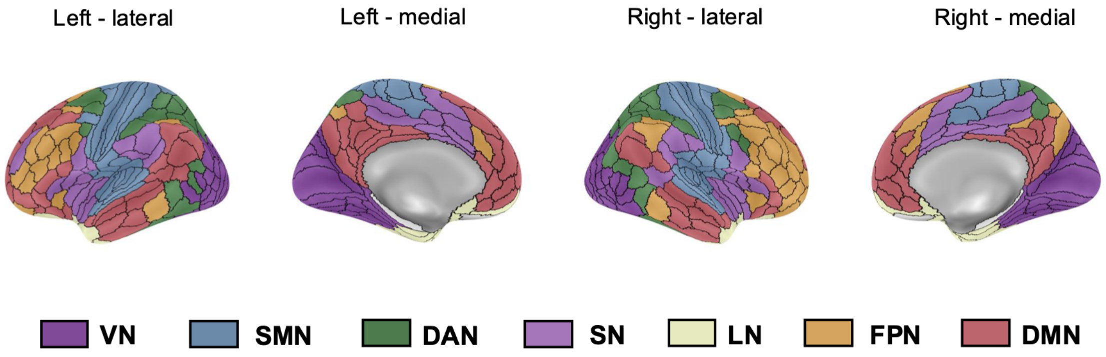


**eFigure 2. Correspondence between HCPex parcels and the seven functional networks.** VN, visual network; SMN, somatomotor network; DAN, dorsal attention network; SN, salience/ventral attention network; LN, limbic network; FPN, frontoparietal control network; DMN, default mode network.

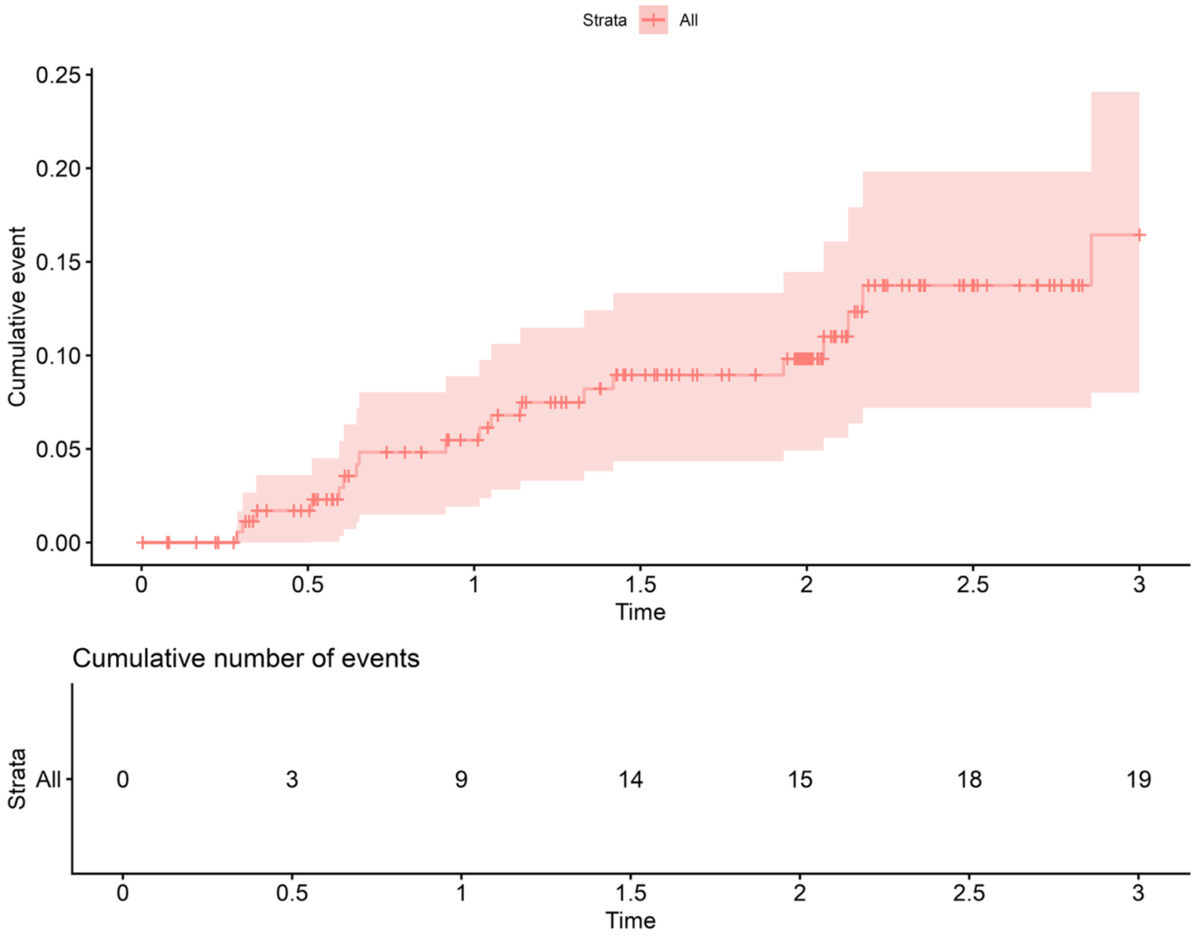


**eFigure 3. Cumulative incidence (Kaplan–Meier failure function) for risk of development of psychosis among CHR-P individuals.** Time on the figure is presented in years. The cumulative incidence of psychosis was 1.7 (95%CI: 0-3.6, 167 individuals still at risk) at 6 months, 5.5 (95%CI: 1.9-8.9, 143 individuals still at risk) at 12 months, 9 (95%CI: 4.3-13.3, 118 individuals still at risk) at 18 months and 9.8 (95%CI: 4.9-14.5, 94 individuals still at risk) at 24 months and 16.4 (95%CI: 8-24.1, 31 individuals still at risk) at 36 months.

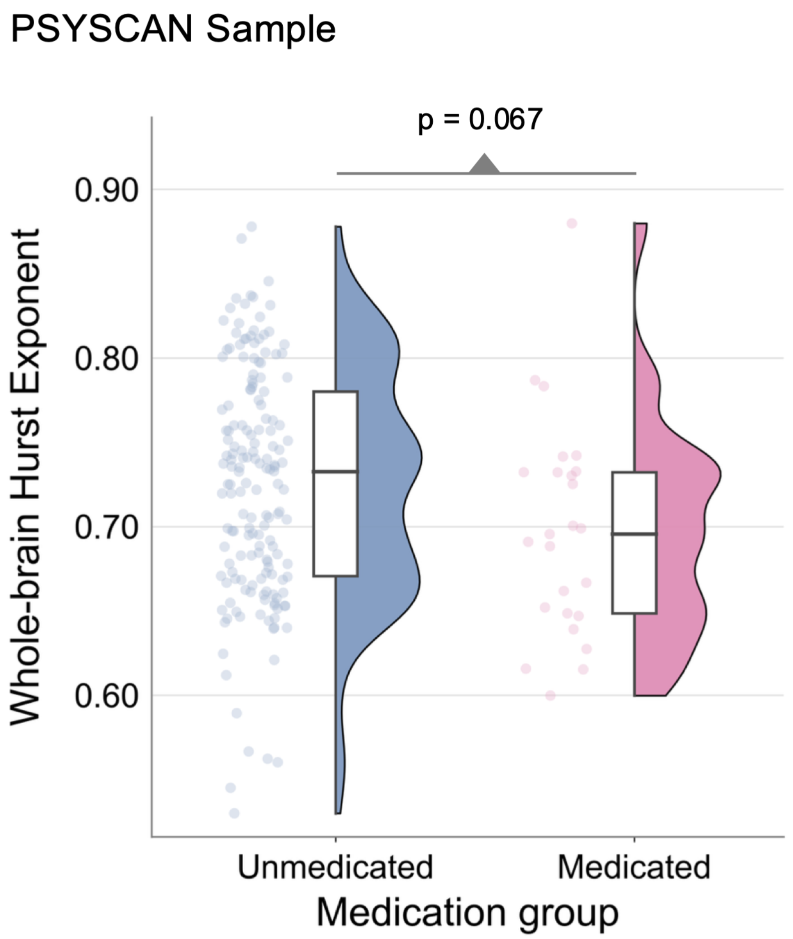


**eFigure 4. Whole-brain Hurst exponent by antipsychotic medication status in PSYSCAN.** Each point represents an individual participant's whole-brain mean Hurst exponent (HE) value, adjusted for age, sex, and mean framewise displacement (MeanFD). Half-violin plots show the distribution of values within each group. In the box plots, the lower and upper edges of the boxes indicate the first and third quartiles, and the horizontal line within each box indicates the median. The vertical whiskers extend to the minimum and maximum observed values.

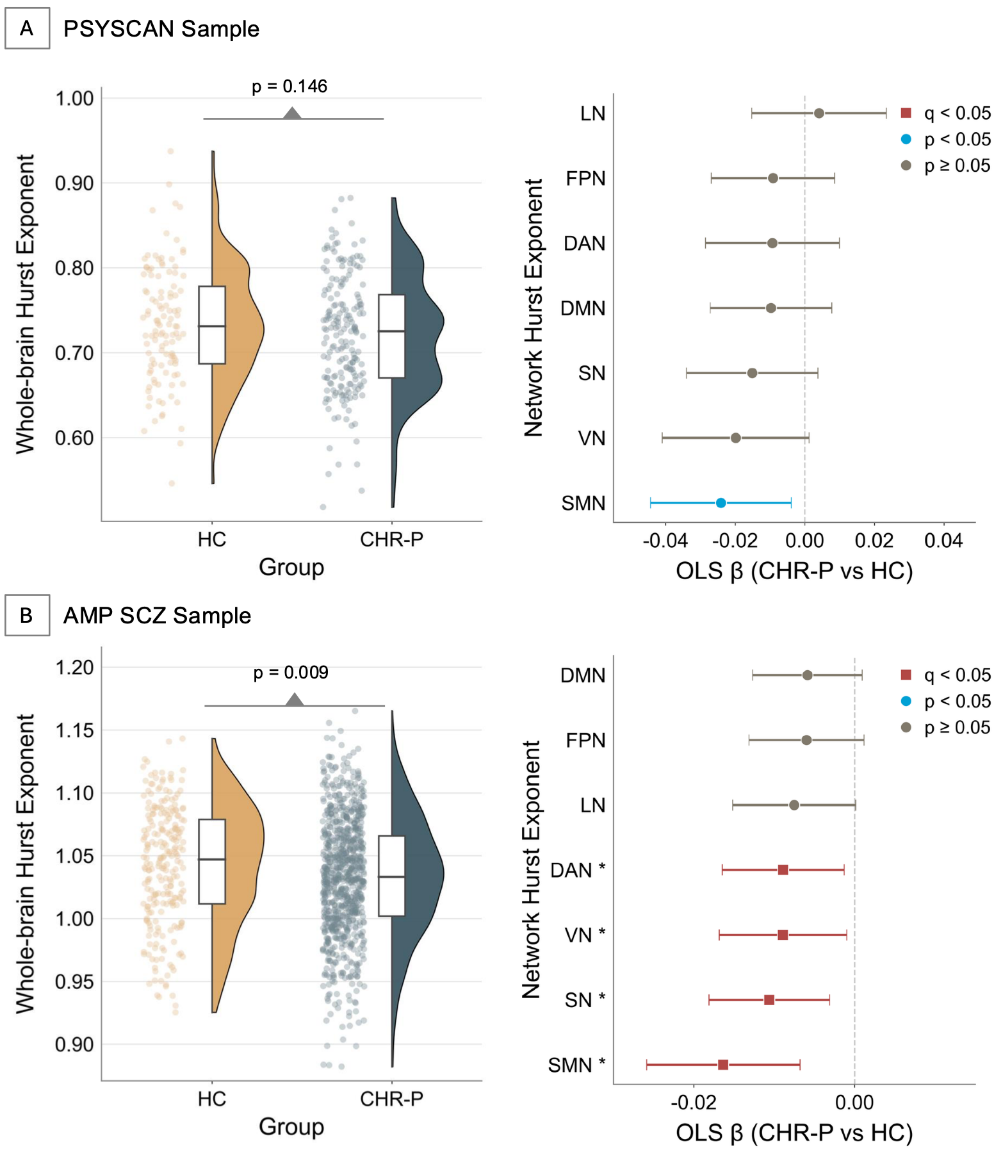


**eFigure 5. Phenotype effects on whole-brain and network-level Hurst exponent after excluding outliers.** A) Whole-brain and network-level Hurst exponent (HE) differences between clinical high risk for psychosis (CHR-P) participants and healthy controls (HCs) in the PSYSCAN sample. B) Corresponding whole-brain and network-level results in the AMP SCZ sample. In the whole-brain plots, each point represents an individual participant's whole-brain mean HE value, adjusted for age, sex, and mean framewise displacement (MeanFD). Half-violin plots show the group-wise distributions, and box plots show the median and interquartile range. In the network-level forest plots, points represent ordinary least squares regression coefficients (OLS β) for the CHR-P versus HC contrast, with horizontal error bars indicating 95% confidence intervals. Red squares indicate effects surviving Benjamini–Hochberg false discovery rate (FDR) correction at $q < 0.05$, blue circles indicate nominally significant effects at $p < 0.05$, and gray circles indicate nonsignificant effects. Asterisks mark networks surviving FDR correction. DAN, dorsal attention network; DMN, default mode network; FPN, frontoparietal control network; LN, limbic network; SMN, somatomotor network; SN, salience/ventral attention network; VN, visual network.

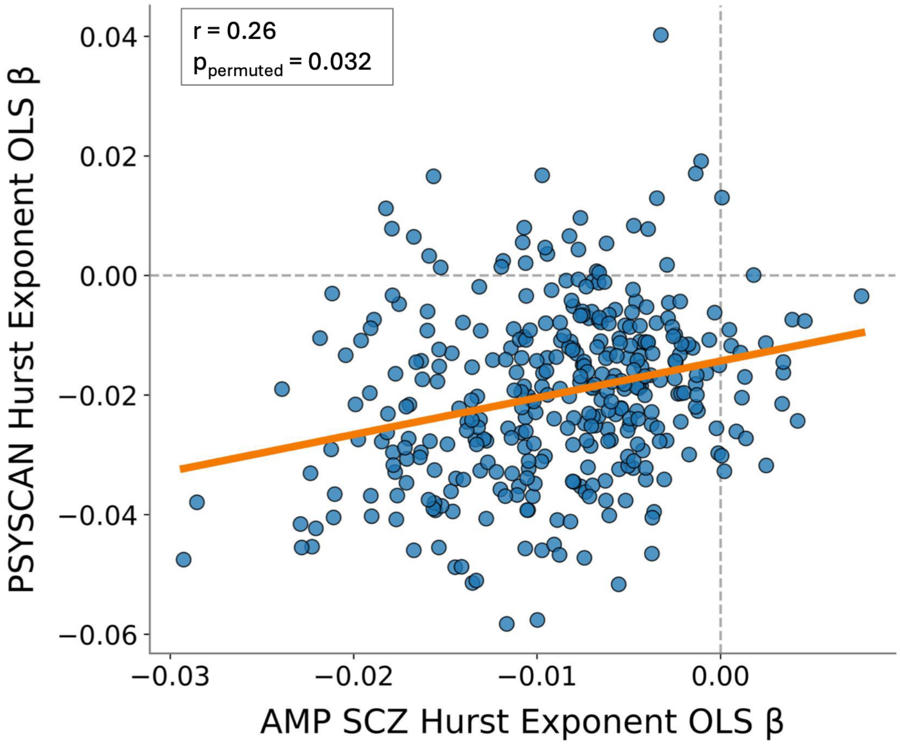


**eFigure 6. Cross-cohort spatial correspondence of phenotype effects on Hurst exponent.** Scatter plot shows the relationship between parcel-wise phenotype effects on Hurst exponent in the AMP SCZ and PSYSCAN samples. Each point represents one cortical parcel. The x- and y-axes show ordinary least squares regression coefficients (OLS β) for the CHR-P versus healthy control contrast in AMP SCZ and PSYSCAN, respectively. The orange line represents the least-squares regression fit, and dashed gray lines indicate zero effects. The Pearson correlation coefficient and permutation-based p-value are shown.

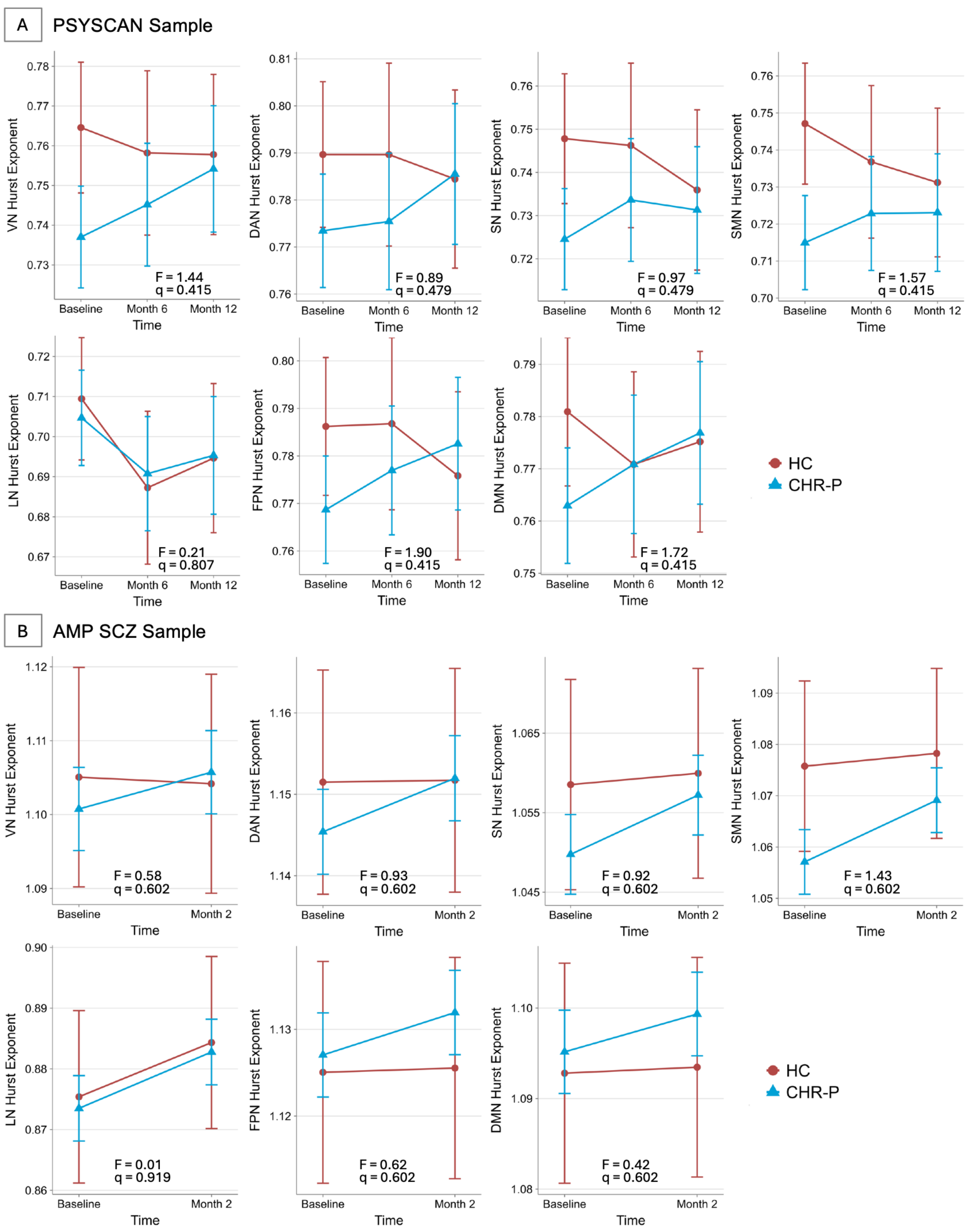


**eFigure 7. Phenotype effects on longitudinal trajectories of network-level Hurst exponent.** A) Longitudinal trajectories of network-level Hurst exponent (HE) in clinical high risk for psychosis participants (CHR-P) and healthy controls (HCs) across seven functional networks in the PSYSCAN sample. B) Corresponding network-level longitudinal trajectories in the AMP SCZ sample. Points represent model-estimated marginal means adjusted for age, sex, and mean framewise displacement (MeanFD), and vertical error bars indicate 95% confidence intervals. Lines connect estimated marginal means across time points. Model statistics indicate the group-by-time interaction effect from linear mixed-effects models, with q-values adjusted using Benjamini–Hochberg false discovery rate correction across the seven networks. VN, visual network; DAN, dorsal attention network; SN, salience/ventral attention network; SMN, somatomotor network; LN, limbic network; FPN, frontoparietal control network; DMN, default mode network.

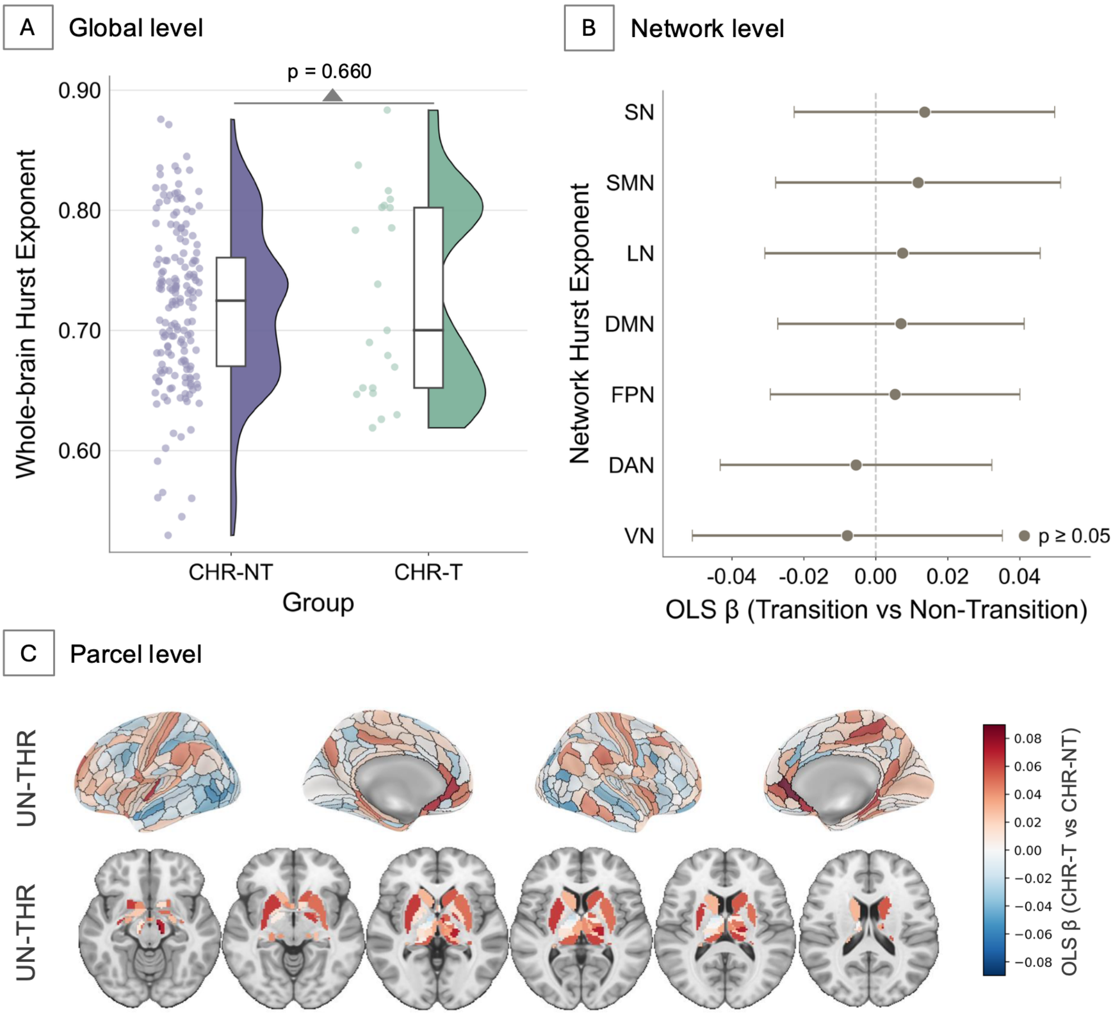


**eFigure 8. Transition-related effects on Hurst exponent in the PSYSCAN sample.** A) Whole-brain Hurst exponent (HE) differences between CHR-P individuals who later transitioned to psychosis (CHR-T) and those who did not transition (CHR-NT) at baseline. B) Network-level HE differences between CHR-T and CHR-NT individuals. C) Parcel-wise cortical and subcortical HE differences between CHR-T and CHR-NT individuals. In the whole-brain plot, each point represents an individual participant's whole-brain mean HE value adjusted for age, sex, and mean framewise displacement (MeanFD). Half-violin plots show the group-wise distributions, and box plots show the median and interquartile range. In the network-level forest plot, points represent ordinary least squares regression coefficients (OLS β) for the CHR-T versus CHR-NT contrast, with horizontal error bars indicating 95% confidence intervals. In the parcel-wise maps, color bars represent OLS β for the CHR-T versus CHR-NT contrast. Blue colors indicate regions with lower HE in CHR-T than in CHR-NT individuals, whereas red colors indicate regions with higher HE in CHR-T than in CHR-NT individuals. Unthresholded maps are shown. DAN, dorsal attention network; DMN, default mode network; FPN, frontoparietal control network; LN, limbic network; SMN, somatomotor network; SN, salience/ventral attention network; VN, visual network.

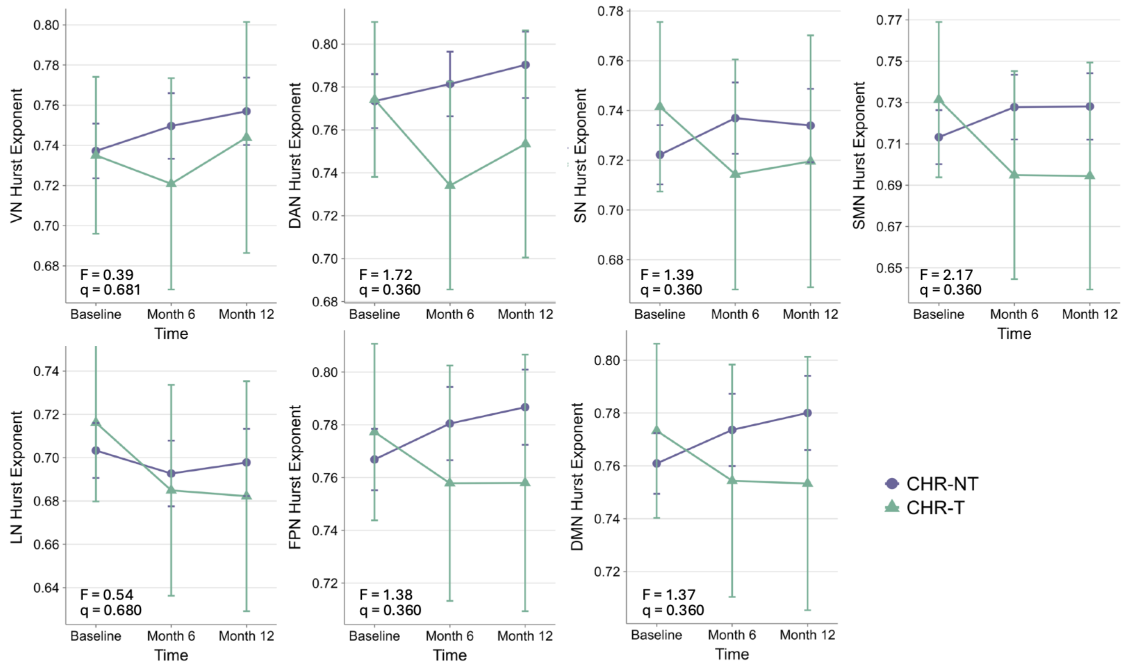


**eFigure 9. Transition-related effects on longitudinal trajectories of network-level Hurst exponent.** Longitudinal trajectories of network-level Hurst exponent (HE) in transition (CHR-T) and non-transition (CHR-NT) participants across seven functional networks in the PSYSCAN sample. Points represent model-estimated marginal means adjusted for age, sex, and mean framewise displacement (MeanFD), and vertical error bars indicate 95% confidence intervals. Lines connect estimated marginal means across time points. Model statistics indicate the group-by-time interaction effect from linear mixed-effects models, with q-values adjusted using Benjamini–Hochberg false discovery rate correction across the seven networks. VN, visual network; DAN, dorsal attention network; SN, salience/ventral attention network; SMN, somatomotor network; LN, limbic network; FPN, frontoparietal control network; DMN, default mode network.

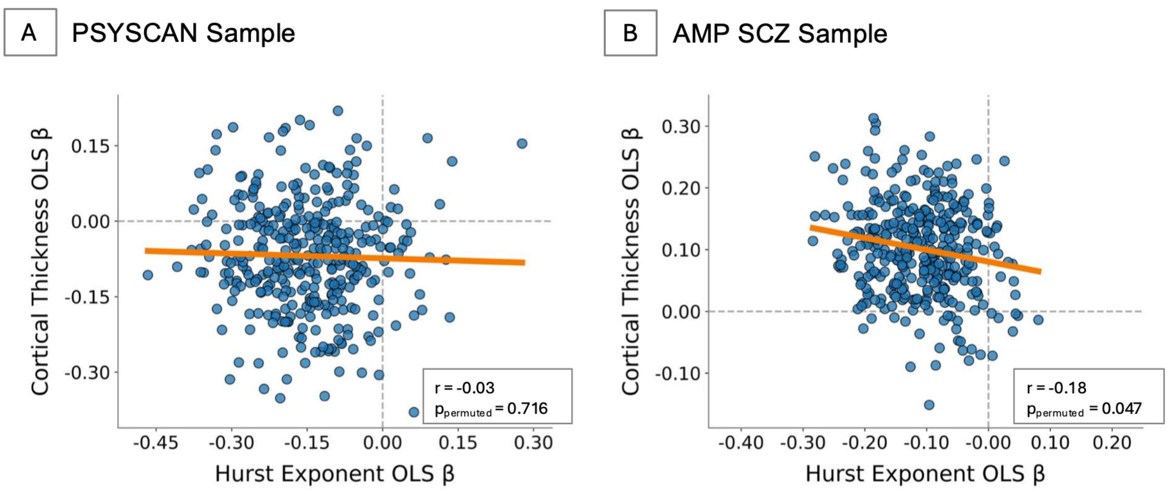


**eFigure 10. Association between phenotype effects on Hurst exponent and cortical thickness across cortical parcels.** Scatter plots show the relationship between parcel-wise phenotype effects on Hurst exponent and cortical thickness in A) the PSYSCAN sample and B)

the AMP SCZ sample. Each point represents one cortical parcel. The x-axis shows ordinary least squares regression coefficients (OLS β) for phenotype effects on Hurst exponent, and the y-axis shows OLS β for phenotype effects on cortical thickness, adjusted for age, sex, and mean framewise displacement (MeanFD). The orange line represents the least-squares regression fit, and dashed gray lines indicate zero effects. Pearson correlation coefficients and permutation-based p-values are shown in each panel. Negative correlations indicate that parcels showing stronger positive phenotype effects on HE tend to show weaker or more negative phenotype effects on cortical thickness, and vice versa.

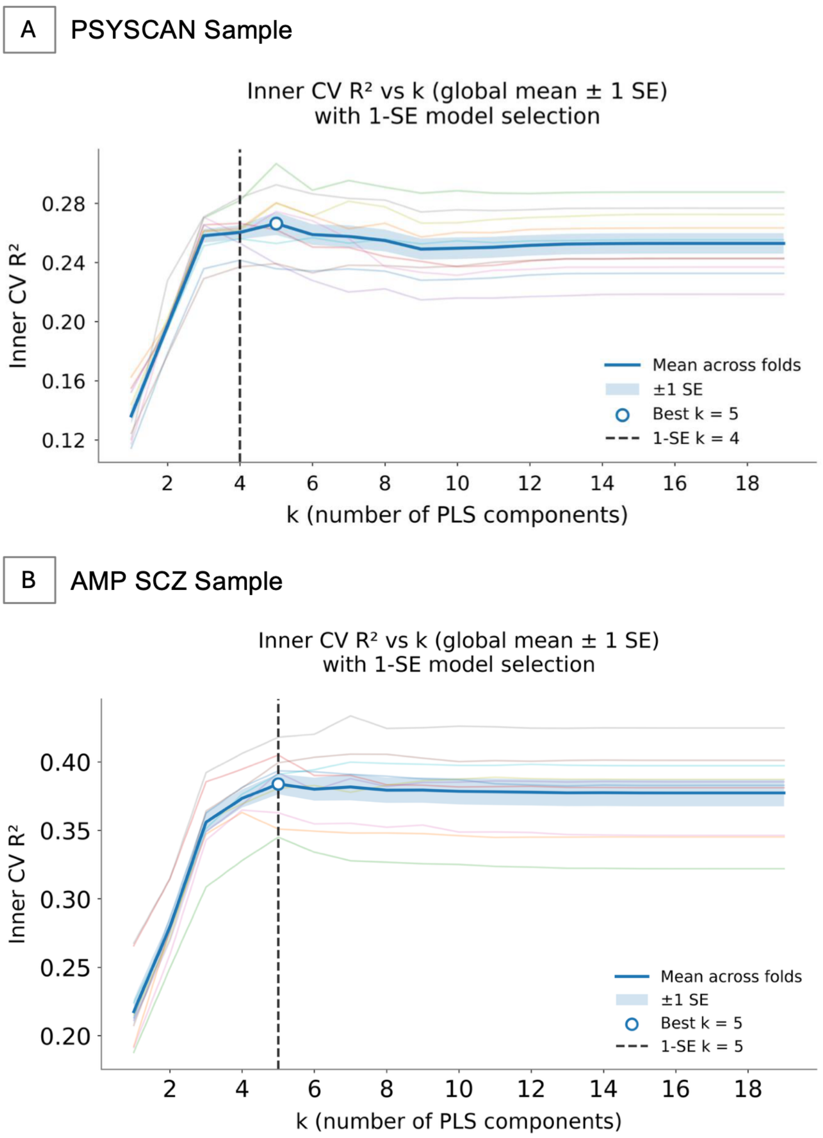


**eFigure 11. Selection of the number of partial least squares components using inner cross-validation.** Plots show inner cross-validation $R^2$ across different numbers of partial least squares components in A) the PSYSCAN sample and B) the AMP SCZ sample. Thin colored lines show individual outer-fold inner cross-validation curves, the solid blue line represents the global mean across folds, and the shaded area indicates ±1 standard error. The open circle marks the number of components with the highest mean inner cross-validation $R^2$, and the dashed vertical line marks the selected number of components based on the one-standard-error rule for the parsimonious model.

### References


1. Tognin S, Van Hell HH, Merritt K, et al. Towards precision medicine in psychosis: benefits and challenges of multimodal multicenter studies—PSYSCAN: translating neuroimaging findings from research into clinical practice. *Schizophr Bull*. 2020;46(2):432-441. doi:10.1093/schbul/sbz067

2. Yung AR, Yuen HP, McGorry PD, et al. Mapping the onset of psychosis: the Comprehensive Assessment of At-Risk Mental States. *Aust N Z J Psychiatry*. 2005;39(11-12):964-971. doi:10.1080/j.1440-1614.2005.01714.x

3. Miller TJ, McGlashan TH, Rosen JL, et al. Prodromal assessment with the structured interview for prodromal syndromes and the scale of prodromal symptoms: predictive validity, interrater reliability, and training to reliability. *Schizophr Bull*. 2003;29(4):703-715. doi:10.1093/oxfordjournals.schbul.a007040

4. Schultze-Lutter F, Ruhrmann S, Picker H, Klosterkötter J. Development and evaluation of the schizophrenia proneness instrument, adult version (SPI-A). *Schizophr Res*. 2006;86:S4-S5. doi:10.1016/S0920-9964(06)70014-7

5. Wannan CMJ, Nelson B, Addington J, et al. Accelerating medicines partnership® schizophrenia (AMP® SCZ): rationale and study design of the largest global prospective cohort study of clinical high risk for psychosis. *Schizophr Bull*. 2024;50(3):496-512. doi:10.1093/schbul/sbae011

6. Woods SW, Parker S, Kerr MJ, et al. Development of the PSYCHS: Positive SYmptoms and Diagnostic Criteria for the CAARMS Harmonized with the SIPS. *Early Interv Psychiatry*. 2024;18(4):255-272. doi:10.1111/eip.13457

7. Harms MP, Cho KIK, Anticevic A, et al. The MR neuroimaging protocol for the Accelerating Medicines Partnership® Schizophrenia Program. *Schizophrenia (Heidelb)*. 2025;11(1):52. doi:10.1038/s41537-025-00581-6

8. Esteban O, Markiewicz CJ, Blair RW, et al. fMRIPrep: a robust preprocessing pipeline for functional MRI. *Nat Methods*. 2019;16(1):111-116. doi:10.1038/s41592-018-0235-4

9. Tustison NJ, Avants BB, Cook PA, et al. N4ITK: improved N3 bias correction. *IEEE Trans Med Imaging*. 2010;29(6):1310-1320. doi:10.1109/TMI.2010.2046908

10. Avants BB, Tustison NJ, Song G, Cook PA, Klein A, Gee JC. A reproducible evaluation of ANTs similarity metric performance in brain image registration. *Neuroimage*. 2011;54(3):2033-2044. doi:10.1016/j.neuroimage.2010.09.025

11. Jenkinson M, Beckmann CF, Behrens TEJ, Woolrich MW, Smith SM. FSL. *Neuroimage*. 2012;62(2):782-790. doi:10.1016/j.neuroimage.2011.09.015

12. Dale AM, Fischl B, Sereno MI. Cortical surface-based analysis. I. Segmentation and surface reconstruction. *Neuroimage*. 1999;9(2):179-194. doi:10.1006/nimg.1998.0395

13. Cox RW. AFNI: software for analysis and visualization of functional magnetic resonance neuroimages. *Comput Biomed Res*. 1996;29(3):162-173. doi:10.1006/cbmr.1996.0014

14. Power JD, Mitra A, Laumann TO, Snyder AZ, Schlaggar BL, Petersen SE. Methods to detect, characterize, and remove motion artifact in resting state fMRI. *Neuroimage*. 2014;84:320-341. doi:10.1016/j.neuroimage.2013.08.048

15. Greve DN, Fischl B. Accurate and robust brain image alignment using boundary-based registration. *Neuroimage*. 2009;48(1):63-72. doi:10.1016/j.neuroimage.2009.06.060

16. Patel AX, Kundu P, Rubinov M, et al. A wavelet method for modeling and despiking motion artifacts from resting-state fMRI time series. *Neuroimage*. 2014;95:287-304. doi:10.1016/j.neuroimage.2014.03.012

17. Greicius MD, Krasnow B, Reiss AL, Menon V. Functional connectivity in the resting brain: A network analysis of the default mode hypothesis. *Proc Natl Acad Sci USA*. 2003;100(1):253-258. doi:10.1073/pnas.0135058100

18. Trakoshis S, Martínez-Cañada P, Rocchi F, et al. Intrinsic excitation-inhibition imbalance affects medial prefrontal cortex differently in autistic men versus women. *eLife*. 2020;9:e55684. doi:10.7554/eLife.55684

19. Uscătescu LC, Hyatt CJ, Dunn J, et al. Using the excitation/inhibition ratio to optimize the classification of autism and schizophrenia. *Transl Psychiatry*. 2025;15(1):234. doi:10.1038/s41398-025-03455-8

20. Fortin JP, Cullen N, Sheline YI, et al. Harmonization of cortical thickness measurements across scanners and sites. *Neuroimage*. 2018;167:104-120. doi:10.1016/j.neuroimage.2017.11.024

21. Huang CC, Rolls ET, Feng J, Lin CP. An extended Human Connectome Project multimodal parcellation atlas of the human cortex and subcortical areas. *Brain Struct Funct*. 2022;227(3):763-778. doi:10.1007/s00429-021-02421-6

22. Yeo BTT, Krienen FM, Sepulcre J, et al. The organization of the human cerebral cortex estimated by intrinsic functional connectivity. *J Neurophysiol*. 2011;106(3):1125-1165. doi:10.1152/jn.00338.2011

23. Kaplan EL, Meier P. Nonparametric estimation from incomplete observations. *Journal of the American Statistical Association*. 1958;53(282):457-481. doi:10.1080/01621459.1958.10501452

24. Lazarus-Barlow WS, Leeming JH. The natural duration of cancer. *BMJ*. 1924;2(3320):266-267. doi:10.1136/bmj.2.3320.266

25. Simon N, Friedman J, Hastie T, Tibshirani R. Regularization paths for cox's proportional hazards model via coordinate descent. *J Stat Softw*. 2011;39(5):1-13. doi:10.18637/jss.v039.i05

26. Fischl B, Dale AM. Measuring the thickness of the human cerebral cortex from magnetic resonance images. *Proc Natl Acad Sci U S A*. 2000;97(20):11050-11055. doi:10.1073/pnas.200033797

27. Hansen JY, Shafiei G, Markello RD, et al. Mapping neurotransmitter systems to the structural and functional organization of the human neocortex. *Nat Neurosci*. 2022;25(11):1569-1581. doi:10.1038/s41593-022-01186-3

28. Markello RD, Hansen JY, Liu ZQ, et al. neuromaps: structural and functional interpretation of brain maps. *Nat Methods*. 2022;19(11):1472-1479. doi:10.1038/s41592-022-01625-w

29. Varma S, Simon R. Bias in error estimation when using cross-validation for model selection. *BMC Bioinformatics*. 2006;7(1):91. doi:10.1186/1471-2105-7-91

30. Hastie T, Tibshirani R, Friedman J. *The Elements of Statistical Learning*. 2nd ed. Springer New York; 2009. doi:10.1007/978-0-387-84858-7
